# The economics of organellar gene loss and endosymbiotic gene transfer

**DOI:** 10.1101/2020.10.01.322487

**Authors:** Steven Kelly

## Abstract

The endosymbiosis of the bacterial progenitors of mitochondrion and the chloroplast are landmark events in the evolution of life on earth. While both organelles have retained substantial proteomic and biochemical complexity, this complexity is not reflected in the content of their genomes. Instead, the organellar genomes encode fewer than 5% of genes found in living relatives of their ancestors. While many of the 95% of missing organellar genes have been discarded, others have been transferred to the host nuclear genome through a process known as endosymbiotic gene transfer. Here we demonstrate that the difference in the per-cell copy number of the organellar and nuclear genomes presents an energetic incentive to the cell to either delete genes or transfer them to the nuclear genome. We show that, for the majority transferred genes, the energy saved by nuclear-transfer exceeds the costs incurred from importing the encoded protein into the organelle where it can provide its function. Finally, we show that the net energy saved by endosymbiotic gene transfer can constitute an appreciable proportion of total cellular energy budgets, and is therefore sufficient to impart a selectable advantage to the cell. Thus, reduced cellular cost and improved energy efficiency likely played a role in the reductive evolution of mitochondrial and chloroplast genomes and the transfer of organellar genes to the nuclear genome.

**Significance statement:** The endosymbioses of the mitochondrion and the chloroplast were each followed by substantial gene loss and transfer of organellar genes to the nuclear genome. Here we show that the high per-cell copy number of these organellar genomes creates an energetic incentive for the cell to discard genes or transfer them to the nuclear genome. Thus, organellar gene loss and endosymbiotic gene transfer can be intrinsically advantageous to the cell.

## Introduction

Endosymbiosis has underpinned two of the most important innovations in the history of life on Earth (Archibald 2015a; Martin, et al. 2015). The endosymbiosis of the alphaproteobacterium that became the mitochondrion led to the emergence and radiation of the eukaryotes (Yang, et al. 1985; Martin and Müller 1998; Roger, et al. 2017), and the endosymbiosis of the cyanobacterium that became the chloroplast first enabled oxygenic photosynthesis in eukaryotes (Martin and Kowallik 1999; Archibald 2015b). The function and evolution of both organelles is inextricably linked with energy metabolism and the evolution of the eukaryotic cell (Lane and Martin 2010; Lane 2014; Booth and Doolittle 2015a, b; Lane and Martin 2015; Lynch and Marinov 2017; Roger, et al. 2017; Lynch and Marinov 2018), and has given rise to the multicellular organisms that constitute the largest fraction of the biomass of the biosphere (Bar-On, et al. 2018).

Following the endosymbioses of the bacterial progenitors of mitochondrion and the chloroplast there was a dramatic reduction in the gene content of the endosymbiont genomes such that they harbour fewer than 5% of the genes found in their free-living bacterial relatives (Gray, et al. 1999; Timmis, et al. 2004; Green 2011). While many of the original endosymbiont genes have been lost (Lynch, et al. 2006; McCutcheon and Moran 2012; Smith and Keeling 2015; Smith 2016), others have been transferred to the host nuclear genome and their products imported back into the organelle where they function - a process known as endosymbiotic gene transfer (Martin, et al. 2002; Brown 2003; Deusch, et al. 2008; Thiergart, et al. 2012; Dagan, et al. 2013). For example, the mitochondrion of humans (Calvo and Mootha 2010) and chloroplasts of plants (Ferro, et al. 2010) each contain more than 1000 proteins yet their genomes encode fewer than 100 genes. Thus, the reduced gene content of organelles is not representative of their molecular, proteomic, or biochemical complexity.

The process of gene loss and endosymbiotic gene transfer is not unique to the evolution of chloroplasts and mitochondria, but has also been observed concomitant with the endosymbioses of bacteria in insects (McCutcheon and Moran 2012; Husnik, et al. 2013) and the endosymbiosis of the cyanobacterium that became the chromatophore in *Paulinella* (Nakayama and Ishida 2009; Nowack, et al. 2010; Reyes-Prieto, et al. 2010; Singer, et al. 2017; Nowack and Weber 2018). In addition, it has been suggested that lateral gene transfers from diverse bacteria into the host nuclear genome may have contributed to the process of organellar genome reduction in a manner that functionally recapitulates endosymbiotic gene transfer i.e. the endosymbiont gene becomes redundant when an orthologous or functionally equivalent gene from another species is transferred to the nuclear genome (Nowack, et al. 2016; Pittis and Gabaldon 2016). Thus, endosymbiont genome reduction in the presence of functional compensation by lateral and/or endosymbiotic gene transfer are recurring themes in the evolution of organellar and endosymbiont genomes.

Given, the importance of endosymbiotic gene transfer (and functionally equivalent lateral complementation) to the evolution of eukaryotic genomes, several hypotheses have been proposed to explain why it occurs (Herrmann 1997; Martin and Herrmann 1998; Daley and Whelan 2005; Reyes-Prieto, et al. 2006; Speijer, et al. 2020). For example, it has been proposed that lateral and endosymbiotic gene transfer protects endosymbiont genes (and the biological functions they provide) from mutational hazard (Allen and Raven 1996; Lynch, et al. 2006; Smith 2016; Speijer, et al. 2020), and that it enables endosymbiont genes that are otherwise trapped in a haploid genome to recombine and thus escape from Muller’s ratchet (Muller 1964; Lynch 1996; Martin and Herrmann 1998; Lynch, et al. 2006; Neiman and Taylor 2009; Smith 2016). It has also been proposed that endosymbiotic gene transfer is an inevitable consequence of a constant stream of endosymbiont genes entering the nucleus (Doolittle 1998), and that transfer to the nuclear genome allows the host cell to gain better control over the replication and function of the organelle (Herrmann 1997) allowing better cellular network integration (Nowack, et al. 2010; Reyes-Prieto 2015). However, mutation rates of organellar genes are often not higher than nuclear genes (Wolfe, et al. 1987; Lynch, et al. 2006; Lynch, et al. 2007; Drouin, et al. 2008; Smith 2015; Smith and Keeling 2015; Smith 2016; Grisdale, et al. 2019) and therefore effective mechanisms for protection against DNA damage in organelles must exist. Similarly, although there is evidence for the action of Muller’s ratchet in mitochondria (Lynch 1996; Neiman and Taylor 2009) chloroplasts appear largely to escape this effect (Wolfe, et al. 1987; Lynch 1997) likely due to gene conversion (Khakhlova and Bock 2006), and thus it does not fully explain why endosymbiotic gene transfer occurred in both lineages. Finally, the nature of the regulatory advantage for having genes reside in the nuclear genome is difficult to quantify, as bacterial gene expression regulation is no-less effective than in eukaryotes and many eukaryotes utilise polycistronic regulation of gene expression (Guiliano and Blaxter 2006; Michaeli 2011; Gordon, et al. 2015; Gallaher, et al. 2021). Thus, it is unclear whether endosymbiotic gene transfer functions simply as rescue from processes that would otherwise lead to gene loss, or whether there may also be an advantage to the cell for transferring an endosymbiont gene to the nuclear genome.

We hypothesised that an advantage for endosymbiotic gene transfer may arise from the difference in the cost to the cell of encoding a gene in the organellar and nuclear genome. This is because each eukaryotic cell typically contains multiple organelles and each organelle typically harbours multiple copies of the organellar genome (Bendich 1987; Cole 2016). The number of organelles in a cell reflects the biochemical requirement of that cell for those organelles, and the high genome copy number per organelle has been proposed to provide protection against DNA damage (Shokolenko, et al. 2009) and to enable the organelle to achieve high protein abundance for genes encoded in the organellar genome (Bendich 1987). Thus, while a diploid eukaryotic cell contains two copies of the nuclear genome, the same cell can contain hundreds to hundreds of thousands of copies of its organellar genomes (Bendich 1987; Cole 2016). For example, endosymbiotic transfer of a 1000 bp gene from the mitochondrion to the nuclear genome in humans, yeast or *Arabidopsis* would save 5,000,000 bp, 200,000 bp or 100,000 bp of DNA per cell, respectively, and an analogous transfer from the chloroplast genome to the nuclear genome in *Arabidopsis* would save 1,500,000 bp of DNA per cell (see Methods for sources of genome copy numbers). As DNA costs energy and cellular resources to biosynthesise (Lynch and Marinov 2015), we hypothesised that if the energy saved by transferring a gene from the organellar genome to the nuclear genome offset the cost of importing the encoded gene products (proteins) back into the organelle then this would provide a direct energetic advantage to the host cell for endosymbiotic gene transfer. Similarly, if a functionally equivalent gene from another species was laterally acquired by the nuclear genome then there would be an analogous energetic advantage to the host cell to utilise the acquired gene and delete the organellar gene.

He we analyse the relative cost of DNA synthesis and protein import over a broad range of plausible parameter spaces for eukaryotic cells that encompasses total cell protein content, organellar fraction (i.e. the fraction of the total number of protein molecules in a cell that are contained within the organelle), organellar genome copy number, organellar protein abundance, organellar protein import cost, organellar protein import efficiency, and protein turnover rate. Through this we reveal that for the vast majority of plausible parameter space for eukaryotic cells it is energetically favourable to the cell to transfer organellar genes to the nuclear genome and re-import the proteins back to the organelle. We show that the interplay between per-cell organellar genome copy number and per-cell organellar protein abundance determines the magnitude of the energy saved such that it is only energy efficient for the cell to retain genes in the organellar genome if they encoding proteins with very high abundance. Through analysis of the energy saved by endosymbiotic gene transfer in the context of total cellular energy budgets, we demonstrate that the net energetic advantage of endosymbiotic gene transfer is a significant proportion of total cell energy budgets and would thus confer a selectable energetic advantage to the cell. Collectively, these results reveal that enhanced energy efficiency has helped to shape the content and evolution of eukaryotic organellar and nuclear genomes.

## Results

### The cost to the cell to encode a gene in the organellar genome is higher than in the nuclear genome

Eukaryotic cells possessing chloroplasts and/or mitochondria typically have a higher copy number of their organellar genomes than their nuclear genomes (Cole 2016). Accordingly, while a typical diploid cell will have two copies of every gene in the nuclear genome, the same cell will have hundreds to hundreds-of-thousands of copies of every organellar encoded gene (Cole 2016). This difference in per-cell genome copy number means that it costs the cell more DNA to encode a gene in the organellar genome than in the nuclear genome. To provide an illustration of this difference in cost we three model eukaryotes were selected with disparate genome sizes and organellar genome content which are representative of the diverse range of values that have been previously reported (Cole 2016). Here, the cost of encoding a gene in a nuclear or organellar genome was considered to be the ATP cost of the chromosome (organellar or nuclear) divided by the number of genes on that chromosome. This consideration was performed to account for differences in organellar and nuclear genomes such as the presence of introns, structural elements (telomeres, centromeres etc), and regulatory elements. We also included the ATP cost of the requisite number of histone proteins contained in nucleosomes to compute the cost of encoding a gene in the nuclear genome. This revealed that the high per cell organellar genome copy number meant that the ATP cost of encoding a gene in the organellar genome is on average one order of magnitude higher than the cost of encoding a gene in the nuclear genome (Figure 1A). This difference in ATP cost is further enhanced if the cost of just the coding sequences (including nucleosomes but excluding introns and non-coding regions) are compared directly (Figure 1B). This latter scenario is more similar to a recent endosymbiotic gene transfer that arrives in the nuclear genome without introns and acquires these over time (Ahmadinejad, et al. 2010). As the three representative organisms shown here span the range of organellar genome copy numbers that have been observed in eukaryotes (Cole 2016), it follows that the ATP cost to the cell of encoding a gene in the organellar genome is generally higher than the cost of encoding the same gene in the nuclear genome in eukaryotes. Consequently, for any organellar gene the cell may be able to save resources by transferring that gene from the organellar genome to the nuclear genome or by acquiring a functionally equivalent gene through lateral gene transfer.

**Figure 1.**
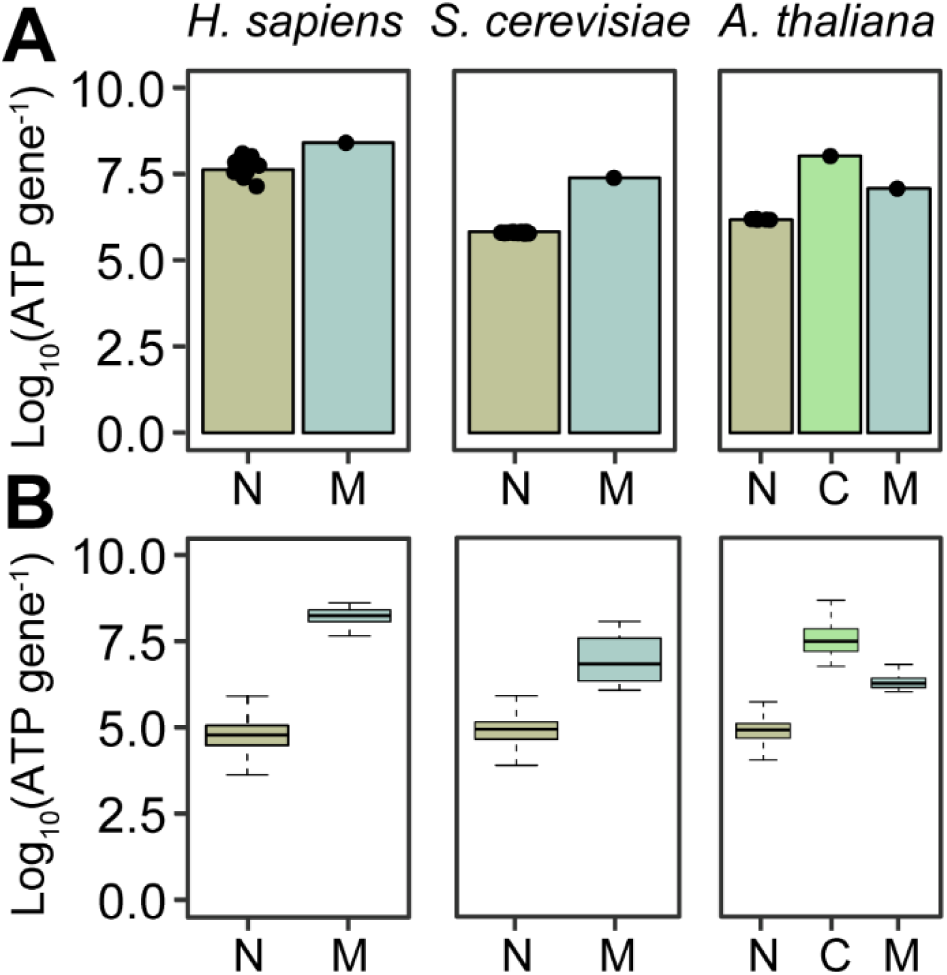
The per-cell biosynthetic cost of nuclear and organellar genes in three representative eukaryotes. **A**) The ATP biosynthesis costs of nuclear (N), chloroplast (C), and mitochondrial (M) genes calculated as the cost of the chromosome divided by the number of genes contained within that chromosome. Nuclear chromosomes include the cost of nucleosomes, organellar chromosomes only included the cost of the DNA. In the case of the nuclear genes the height of bar depicts the mean cost of all nuclear chromosomes with individual points showing all chromosomes overlaid on top the bar plots. **B**) The ATP biosynthesis cost of just the coding sequences of the genes. In both A and B, the costs were computed assuming a diploid nuclear genome, a per-cell mitochondrial genome copy number of 5000, 200 and 100 for the in *H. sapiens*, *S. cerevisiae* and *A. thaliana*, respectively, and a per cell chloroplast genome copy number of 1500 in *A. thaliana*.

### The energy saved by encoding a gene in the nuclear genome instead of the organellar genome is sufficient to offset the cost of organellar protein import

Although it is cheaper for the cell to encode a gene in the nuclear genome than the organellar genome, this direct cost comparison only considers the cost of DNA (and its associated proteins) and does not account for the additional cost that would be incurred should the product of a nuclear encoded gene be required to function in an organelle. Such nuclear-encoded organelle-targeted proteins incur additional energetic costs to be translocated across the organellar membranes. Accordingly, to assess whether it is cheaper for the cell to encode an organelle-targeted protein in the nuclear or organellar genome it is necessary to consider both the abundance of the encoded protein and the energetic cost of organellar protein import. Estimates for the energetic cost of mitochondrial or chloroplast protein import vary over two orders of magnitude from ∼0.05 ATP per amino acid to 5 ATP per amino acid (Mokranjac and Neupert 2008; Shi and Theg 2013; Backes and Herrmann 2017). Thus, for the purposes of this study the full range of estimates was considered and the range of conditions under which it is more energetically favourable to encode a gene in the organellar or nuclear genome was assessed. This analysis revealed that the higher the copy number of the organellar genome, the more energy that is saved by encoding the gene in the nuclear genome and thus the more protein that can be imported into the organelle while still reducing the overall energetic cost of the cell (Figure 2A). As the per-cell gene copy number is the same for each gene encoded on the organellar genome, the possible energetic advantage to the cell arising from endosymbiotic gene transfer will vary between genes as a function of the required abundance of each encoded gene product. Furthermore, if the cell can function without the encoded gene product, then as organellar genome copy number increases the energetic incentive to discard the gene also increases. Thus, high organellar genome copy numbers provide an energetic incentive to either delete genes from the organellar genome or transfer them to the nuclear genome.

**Figure 2.**
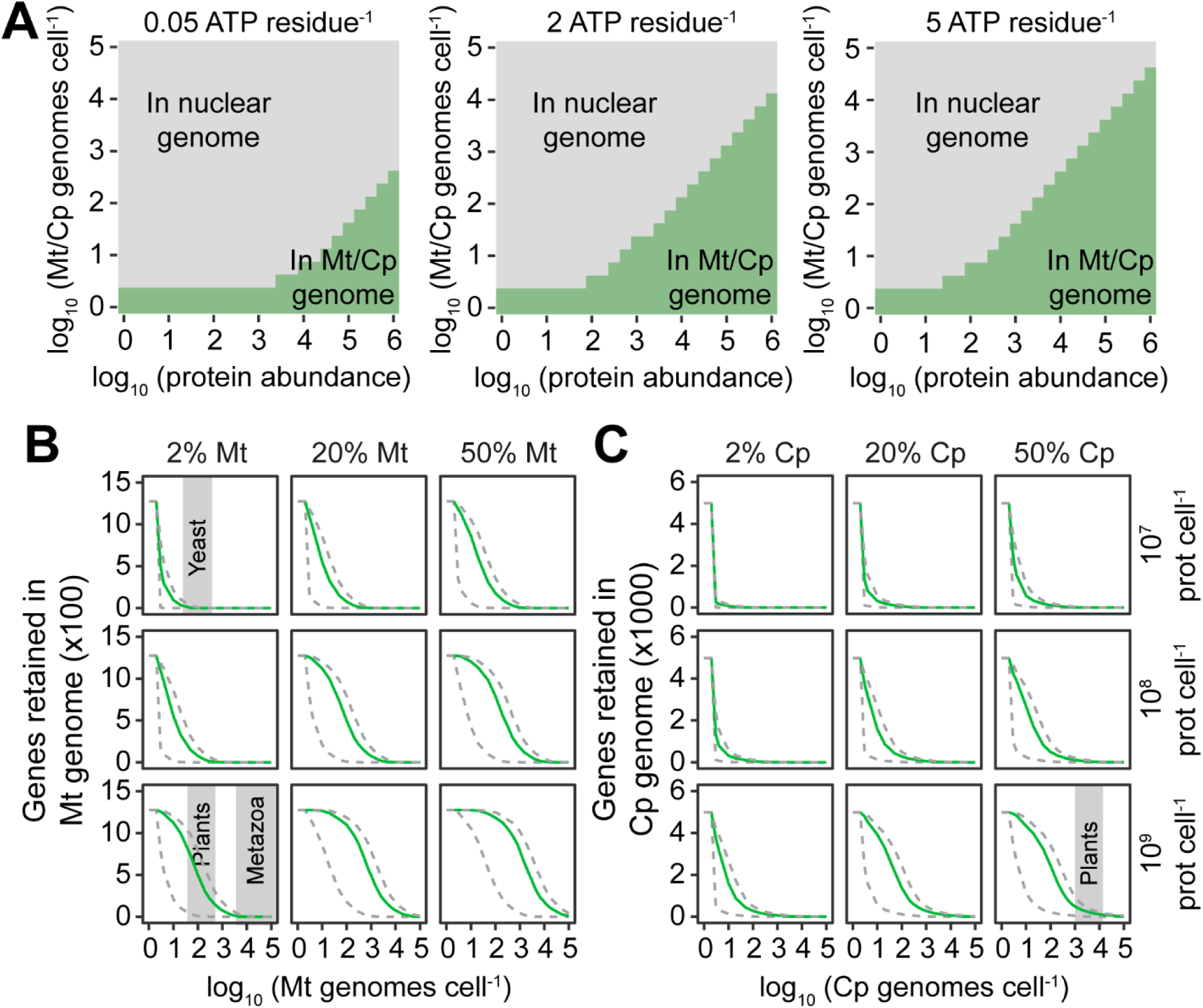
The minimum cost location to the cell of organellar genes encoding an organellar localised protein. **A**) The minimum cost location of an organellar gene for a range of per-protein import costs, organellar genome copy numbers, and encoded protein abundance. The modelled per-residue protein import cost is shown above each plot. The grey shaded fractions of the plots indicate the regions of parameter space where it is more energetically favourable to the cell to encode an organellar gene in the nuclear genome and import the requisite amount of protein. The green shaded fractions of the plots indicate the regions of parameter space where it is more energetically favourable to the cell to encode the gene in the organellar genome. **B**) The number of genes in the alphaproteobacterial (mitochondrial) genome for which it is more energetically favourable to the cell for the gene to be retained in the organellar genome. Green lines assume a per-residue protein import cost of 2 ATP per amino acid. Grey dashed lines indicate lower and upper cost bounds of 0.05 ATP and 5 ATP per residue respectively. **C**) As in B but for the cyanobacterial (chloroplast) genome. Grey shaded areas on plots are provided to indicate the organellar genome copy numbers of yeast, metazoan and plant cells. Cp: chloroplast. Mt: mitochondrion.

Given that the magnitude of the energetic advantage of endosymbiotic gene transfer is dependent on protein abundance, we sought to simulate the endosymbiotic genome reduction that would occur using realistic models of pre-mitochondrial and pre-chloroplast organellar progenitors. Here, the complete genomes with measured protein abundances for an alphaproteobacterium (*Bartonella henselae*) and a cyanobacterium (*Microcystis aeruginosa*) were chosen as models for the mitochondrial and chloroplast progenitors, respectively. In addition, a range of host cell size (i.e. host cell protein content) was considered such that it encompassed the majority of diversity exhibited by extant eukaryotes (Milo 2013) and would thus likely also encompass the size range of the host cell that originally engulfed the organellar progenitors. This range extended from a small unicellular yeast-like cell (10^7^ proteins) to a large metazoan/plant cell (10^9^ proteins). Each of these cell types were then considered to allocate a realistic range of total cellular protein to mitochondria/chloroplasts representative of values observed in extant eukaryotic cells (Supplemental Table S1). For each set of conditions in this comprehensive parameter space, the energy liberated or incurred by endosymbiotic gene transfer was calculated for each organellar gene given its measured protein abundance (Wang, et al. 2015) and a realistic range of protein import costs (including the total biosynthetic cost of the protein import machinery, See Methods). This revealed that for a broad range of estimates of cell size, organellar genome copy number, and organellar fraction (i.e. the fraction of the total number of protein molecules in a cell that are contained within the organelle) it is energetically favourable to the cell to transfer the majority of organellar genes to the nuclear genome and re-import the proteins back to the organelle (Figure 2B and 2C). Only the proteins with the highest abundance, and thus which incur the largest import cost, are energetically favourable to be retained in the organellar genomes. This phenomenon was also observed even if extreme costs for protein import ten times those that have been measured are considered (Supplemental Figure S1). Thus, it is more energy efficient for a eukaryotic cell to position the majority of genes that encode organellar targeted proteins in the nuclear genome.

The above analysis assumed that the total pool of cellular protein was replaced with each cell doubling. This assumption is consistent with observations that protein turn-over in eukaryotes (as in bacteria) is primarily mediated by dilution due to cell division (Boisvert, et al. 2012; Gawron, et al. 2016; Martin-Perez and Villén 2017), i.e. the vast majority of proteins have half-lives that are longer in duration than the doubling time of the cell, and thus protein turn-over occurs through replicative dilution. However, a small population of proteins are turned-over more than once per cell division cycle (Boisvert, et al. 2012; Gawron, et al. 2016; Martin-Perez and Villén 2017), and in multicellular organisms there can be populations of cells with low or negligible rate of cell division resulting in a higher rate of protein turn-over per cell division. Thus, to determine the impact of enhanced rates of protein turn-over relative to cell doubling the analysis above was repeated while increasing the rate of protein turn-over from once per cell division cycle (i.e. dividing cells) to 50 times per cell division cycle (i.e. a long-lived or non-dividing cell). Increasing the rate of protein turn-over increases the total amount of protein that must be imported into the organelle (akin to an increase in absolute abundance of that protein) and thus leads to an increase in the number of proteins for which it is energetically favourable to retain their corresponding genes in the organellar genomes (Figure 3A and B). However, even if it is assumed that the total pool of each organellar protein is turned over 50 times per cell division cycle, it is still more energetically favourable to transfer the majority of organellar genes to the nuclear genome when organellar genome copy number is high (Figure 3A and B). Thus, in both dividing cells and in cells with higher rates of protein turn-over relative to cell division it is more energetically favourable to encode the majority of organellar targeted proteins in the nuclear genome.

**Figure 3.**
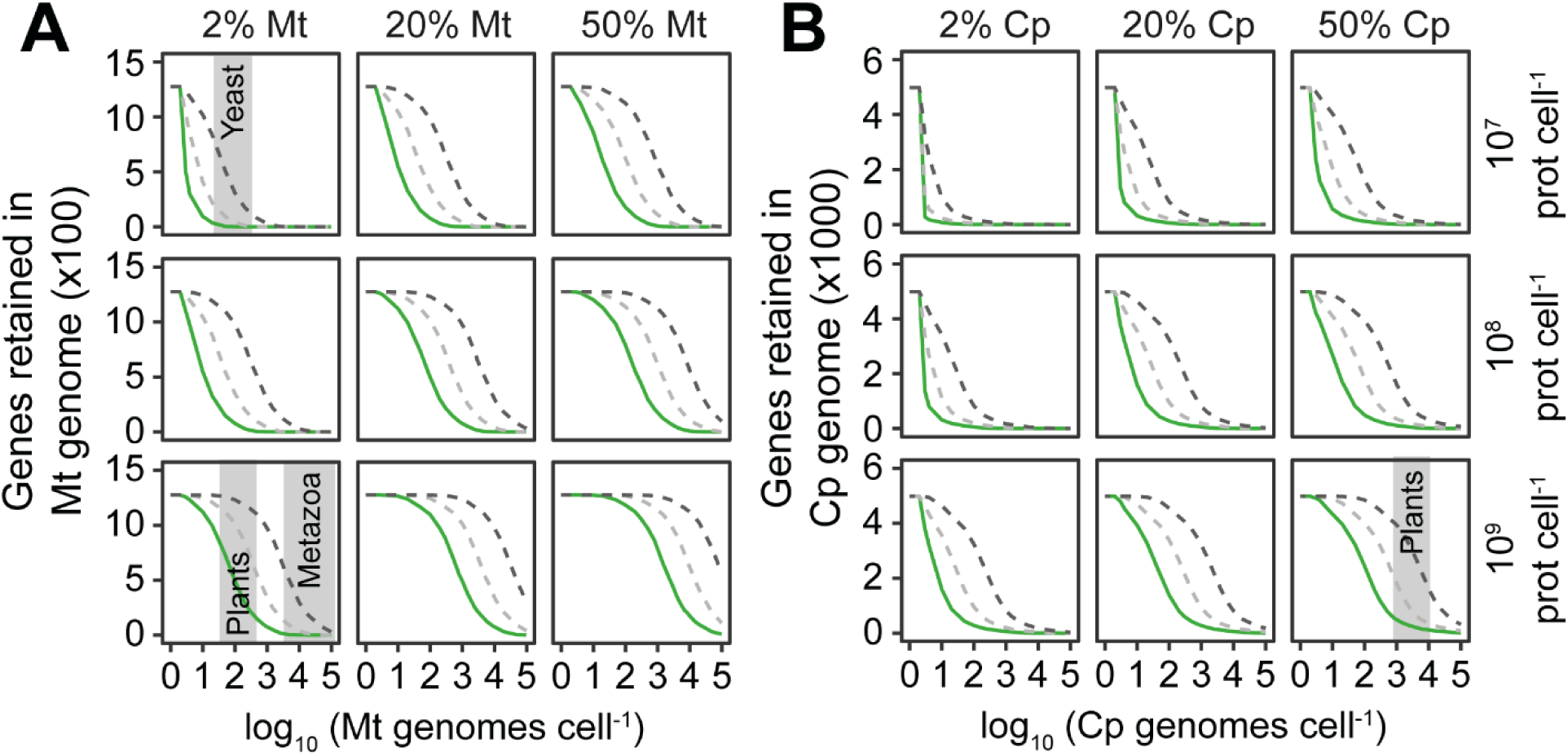
The impact of protein turnover on the energetic favourability or organellar gene retention. **A)** The number of genes in the alphaproteobacterial (mitochondrial) genome for which it is more energetically favourable to the cell for the gene to be retained in the organellar genome. **C**) As in B but for the cyanobacterial (chloroplast) genome. All lines assume a per-residue protein import cost of 2 ATP per amino acid. Green lines assume that protein turnover is mediated by dilution due to cell division. Light grey dashed lines assume that the complete pool of organellar proteins at the requisite abundance are replaced 5 times per cell doubling. Dark grey dashed lines assume that the complete pool of organellar proteins at the requisite abundance are replaced 50 times per cell doubling. Grey shaded areas on plots are provided for illustrative purposes to indicate the organellar genome copy numbers of yeast, metazoan and plant cells. Cp: chloroplast. Mt: mitochondrion.

### Protein abundance can explain a significant proportion of variance in loss, retention and, transfer of organellar genes

The analyses above predict that the proteins with the highest abundance, and thus those which incur the higher import cost, are those that are more likely to be retained in an organellar genome. While it is unknown how the abundance of proteins in organelles has changed throughout the evolution of the eukaryotes, it is possible to estimate what the profile of protein abundances may have looked like during the initial stages of this process by examining protein abundance in extant bacterial relatives of organelles (Wang, et al. 2015). Using these estimates it is thus possible to ask whether those genes that are retained in the organellar genome are those that encode proteins with higher abundance than those that are lost or transferred to the nuclear genome. This revealed that the abundance of the cohorts of proteins whose genes are retained in the chloroplast (Figure 4A) and mitochondrial (Figure 4B) genomes of *Arabidopsis thaliana,* and the mitochondrial genome of *Saccharomyces cerevisiae* (Figure 4C) are significantly higher than the abundance of the cohorts of proteins that were either lost or transferred to the respective nuclear genomes. The abundance of the cohort of proteins whose genes are retained in the mitochondrial genome of *Homo sapiens* were not significantly different to those that have been lost or transferred to the nuclear genome (Figure 4D). To assess whether or not this elevated protein abundance was a general phenomenon, the full set of complete plastid and mitochondrial genomes were downloaded from NCBI and the sets of genes present or absent from these genomes were analysed. Here, the corresponding nuclear genomes were not available so it was not possible to separately assess the abundance proteins encoded by lost or putatively transferred genes, and thus they were analysed together. This analysis revealed that the abundance of proteins encoded by genes found in extant plastid (Figure 4E) or mitochondrial (Figure 4F) genomes in eukaryotes was significantly higher than those that have been lost or transferred to the nuclear genome. Thus, across all eukaryotes the abundance of proteins encoded by genes retained in organellar genomes is higher than the abundance of proteins encoded by genes that were either lost or transferred to the nuclear genome.

**Figure 4.**
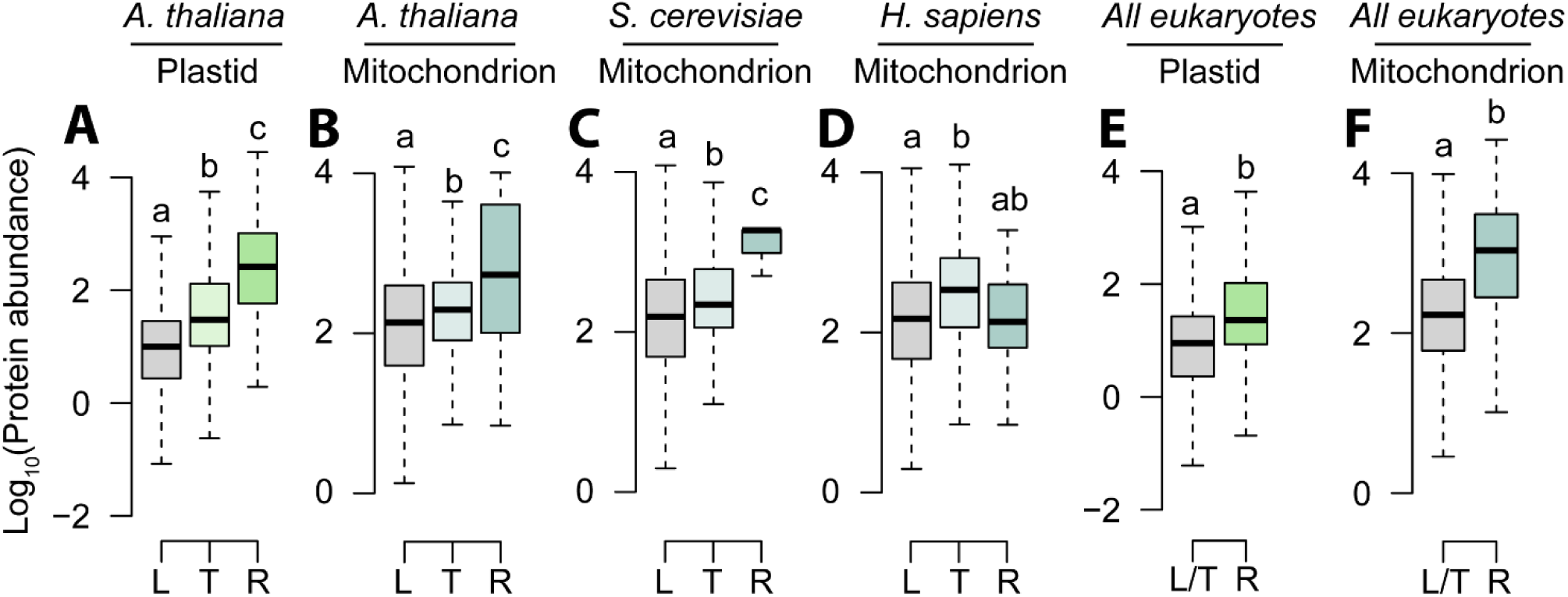
The abundance of proteins encoded by genes that have been lost, transferred to the nucleus or retained in the organellar genome. **A**) The abundance of proteins in the cyanobacterium *Microcystis aeruginosa* categorised according to whether their encoding genes have been lost, transferred to the *Arabidopsis thaliana* nuclear genome, or retained in the *Arabidopsis thaliana* chloroplast genome. **B**) The abundance of proteins in the alphaproteobacterium *Bartonella henselae* categorised according to whether their encoding genes have been lost, transferred to the *Arabidopsis thaliana* nuclear genome, or retained in the *Arabidopsis thaliana* mitochondrial genome. **C** and **D**, as in B but for *Saccharomyces cerevisiae* and *Homo sapiens*, respectively. **E**) as in A, but for all plastid genomes on NCBI. **F**) as in B, but for all mitochondrial genomes on NCBI. L: lost. T: Transferred to nuclear genome. R: Retained in organellar genome. Letters above boxplots indicate whether there were significant differences between the means of different groups (p < 0.05) in the results of a one-way ANOVA with Tukey test for multiple comparisons.

### The energy saved by gene loss or endosymbiotic gene transfer is sufficient to produce a selectable advantage for the majority of genes

Although gene loss or endosymbiotic gene transfer can save energy, the question arises as to whether this energy saving would be sufficient to confer a selectable advantage for the cell. To estimate this, the energy liberated by endosymbiotic gene transfer of each gene encoded in the ancestral pre-organellar genomes was evaluated as a proportion of the total energy required to replicate the cell. As above, this analysis was conducted for a broad range of host cell size, organellar fraction, endosymbiont/organellar genome copy number, and protein import cost that is representative of a broad range of eukaryotic cells (Figure 5A and B, Supplemental Figures S2 – S7, Supplemental Table S2). This revealed that for even modest per-cell endosymbiont genome copy numbers (∼100 copies per cell), the proportion of the total cell energy budget that could be saved for an individual gene transfer event (or equivalent functional lateral complementation) is sufficient that it would confer a selectable advantage. If the energetic advantage is considered to be a direct fitness advantage then the selection coefficients for the transfer of the majority of individual endosymbiont genes are ∼1×10^-5^ (Figure 5, Supplemental Figures S2 – S7). This ∼1,000 times stronger than the selection coefficient acting against disfavoured synonymous codons (Hartl, et al. 1994). Moreover, for high per-cell endosymbiont genome copy numbers (∼1000 genome copies per cell) these selection coefficients are proportionally larger (∼1 × 10^-4^), equivalent to approximately 1/10^th^ the strength of selection that caused the allele conferring lactose tolerance to rapidly sweep through human populations in ∼500 generations (Bersaglieri, et al. 2004). In contrast, selection coefficients for retention of genes in the organellar genome generally only occur when organellar genome copy numbers are low, and/or when large proportions of cellular resources are invested in organelle (Figure 5A and B, Supplemental Figures S2 – S7). Thus, over a broad range of host cell sizes, organellar genome copy numbers, organellar fractions, and per-protein ATP import costs, endosymbiotic gene transfer of the majority of genes is sufficiently energetically advantageous that any such transfer events, if they occurred, would confer an energetic advantage to the cell and have the potential to rapidly reach fixation (Supplemental Figure S8). Thus, endosymbiotic gene transfer of the majority of organellar genes is advantageous to eukaryotic cells.

**Figure 5.**
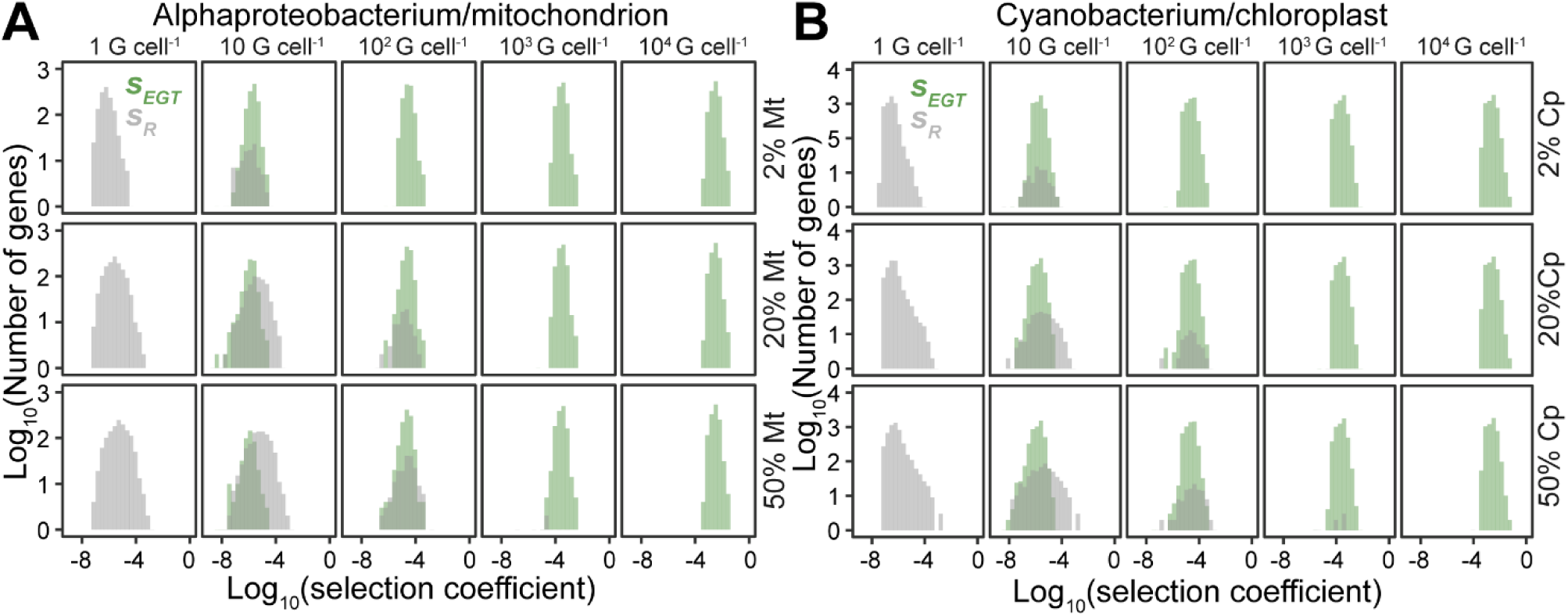
Selection coefficients for retention (*S_R_,* grey) or endosymbiotic gene transfer (*S_EGT_*, green) of all genes encoded in the example alphaproteobacterial and cyanobacterial genomes. Coefficients were computed accounting for protein abundance, host cell organellar fraction, organellar genome copy number per cell, and host cell energy consumption. Plots shown are for a simulated host cell comprising 1 × 10^7^ proteins and a protein import cost of 2 ATP per residue, plots for other host cell protein contents and protein import costs are provided in Supplemental Figures S4-S9. **A**) Selection coefficients of all genes encoded in the alphaproteobacterium genome. **B**) Selection coefficients for all genes encoded in the cyanobacterial genome. *S_R_* and *S_EGT_* have opposite signs (see methods). To simplify the display and enable direct comparison, the absolute value of the selection coefficients of each gene are plotted and green shading is used to indicate genes in the *S_EGT_* fraction and grey shading indicates genes int the *S_R_* fraction of the genome. Mt, mitochondrion. Cp, chloroplast. G, genomes.

## Discussion

The endosymbiosis of the bacterial progenitors of the mitochondrion and the chloroplast are landmark events in the evolution eukaryotes. Following these endosymbioses there was a dramatic reduction in the gene content of the organellar genomes such that they now harbour fewer than 5% of the genes found in their free-living bacterial relatives. Some of these genes have been discarded, but many have been transferred to the nuclear genome and their products (proteins) are imported back into the organelle where they function. The reason why these organelles have transferred their genes to the nucleus is a long-standing unanswered question in evolutionary biology. Here, we show, through extensive simulation of plausible parameter spaces for eukaryotic cells, that there are energy incentives for gene loss and for endosymbiotic gene transfer from organellar genomes. We show that these energy incentives are dependent on the abundance of the encoded gene product, with a trade-off between per-cell organellar genome copy number and protein abundance determining magnitude and direction of the energy incentive. We further show that these energy incentives can be sufficient to produce a selectable advantage to the host cell for both endosymbiotic gene transfer and for retention of genes in the organellar genomes. Thus, the economics of protein production and transport plays a role in determining whether genes are lost, retained or transferred from organellar genomes.

Although this study reveals that the energy efficiency of protein production can provide driver for the location of an organellar gene, it is not proposed that it is the only factor that influences this process. Instead, a large cohort of factors including the requirement for organellar mediated RNA editing, protein chaperones, protein folding, post-translational modifications, escaping mutation hazard, Muller’s rachet, enhanced nuclear control, the requirement for redox regulation of gene expression, and drift will act antagonistically or synergistically with energetic incentives described here to influence the set of genes that are retained in, lost, or transferred from, the organellar genomes. The study presented here simply reveals that energy efficiency is a previously overlooked factor that has likely played a role in shaping the evolution organellar/nuclear genomes. Moreover, the change in cost to the cell provides a simple mechanistic basis for selection to act with or against Doolittle’s “You are what you eat” ratchet for endosymbiotic gene transfer (Doolittle 1998).

It is noteworthy in these contexts, that if the protein encoded by the endosymbiont gene can provide its function outside of the endosymbiont (e.g. by catalysing a reaction that could occur equally well in the cytosol of the host as in the endosymbiont) then the energetic advantage of gene transfer to the nuclear genome is further enhanced, as the cost of protein import is not incurred. Similarly, although gene loss has been proposed to be mediated predominantly by mutation pressure and drift (Lynch, et al. 2006), the elevated per-cell endosymbiont genome copy number also provides a substantial energetic reward to the host cell for complete gene loss as neither the costs of encoding the gene or producing its product are incurred. Thus, high organellar genome copy number provides an energetic incentive for the cell to delete endosymbiont genes or transfer them to the nuclear genome.

While the analysis presented here focussed on the energetic cost measured in ATP so that the cost of protein import and the cost of biosynthesis of DNA could be evaluated on a common basis, endosymbiotic gene transfer also results in changes in the elemental requirements of a cell. Specifically, as the monophosphate nucleotides that constitute DNA are composed of carbon (A = 10, C = 9, G = 10, T = 10), nitrogen (A = 5, C = 3, G = 5, T = 2), and phosphorous (A = 1, C = 1, G = 1, T = 1) atoms, endosymbiotic gene transfer can also result in substantial savings of these resources (Supplemental Figure S9). Thus, if organisms encounter carbon, nitrogen or phosphorous limitation in their diet and environment then the advantage of endosymbiotic gene transfer to the cell will be further enhanced.

The analysis presented here shows that for a broad range of cell sizes and resource allocations that endosymbiotic gene transfer of the majority of organellar genes is energetically favourable and thus advantageous to the cell. However, it also showed that retention of genes in the organellar genomes is energetically favourable under conditions where the encoded organellar protein is required in very high abundance and/or the copy number of the organellar genome is low. Other interlinked competing factors that influence the energetically optimal location of a gene are shown in Figure 6. Each of these factors interact to influence the cost to the cell for encoding a gene in the nuclear or organellar genome. This is important, as while we do not know precisely what the cells that engulfed the progenitors of the mitochondrion or the chloroplast looked like (as only extant derivatives survive), it is safe to assume that cell size and investment in organelles has altered since these primary endosymbioses first occurred. Accordingly, the selective advantage (or disadvantage) of transfer of any given gene is transient and will have varied during the radiation of the eukaryotes as factors such as cell size and organellar volume evolved and changed in disparate eukaryotic lineages. This coupled with the lack of an organellar protein export system (i.e. from the organelle to the host cytosol) and the presence (and acquisition) of introns in nuclear encoded genes (Rogozin, et al. 2012) means that it is more difficult for endosymbiotic gene transfer to operate in the reverse direction (i.e. from the nucleus to organelle). Similarly, eukaryotic cells can typically tolerate the loss of one or more chloroplasts (Zhuang and Jiang 2019) or mitochondria (Ding and Yin 2012) from a cell without the concomitant death of the cell, the disruption of these organelles is thought to be a major route through which DNA from organelles enters the nucleus and can thus be incorporated into the nuclear genome. The converse process (i.e. the loss of the nucleus) is terminal to the cell and is thought to be a major reason why endosymbiotic gene transfer operates in one direction only. Collectively, these factors would create a ratchet-like effect trapping genes in the nuclear genome even if subsequent changes in cell size and organellar fraction means that it became energetically advantageous to return the gene to the organelle later in evolution. Thus, current organellar and nuclear gene contents predominantly reflect past pressures to delete organellar genes or transfer them to the nuclear genome.

**Figure 6.**
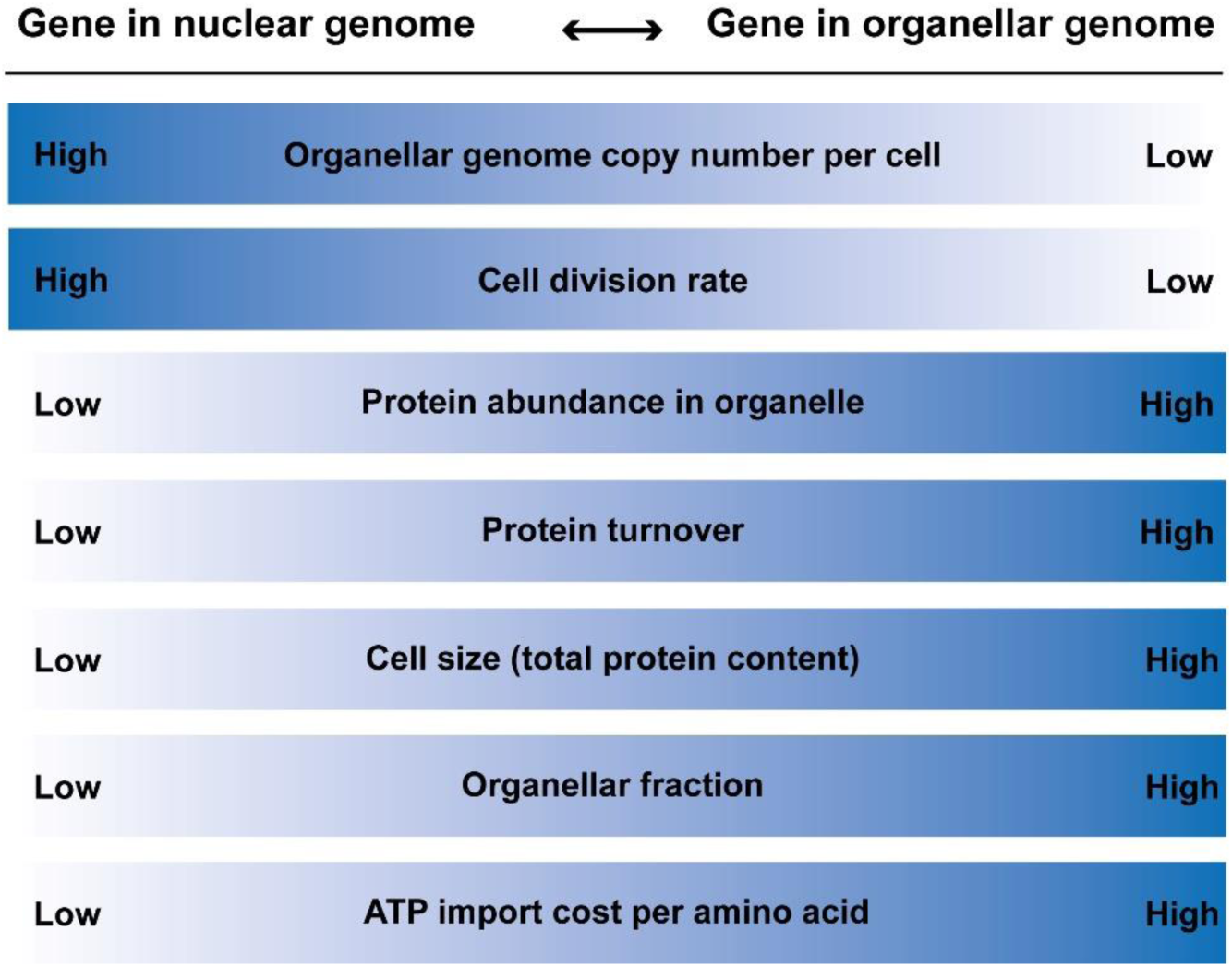
The competing factors that influence the energetically optimal location of a gene encoding an organellar targeted protein. Many of these factors are linked (e.g. protein abundance in organelle and organellar fraction, or cell division rate and protein turn-over) and are provided here for completion.

Endosymbiotic gene loss and gene transfer is a recurring theme in the evolution of the eukaryotic tree of life. The discovery that endosymbiotic gene transfer (or equivalent functional lateral complementation) can provide an energetic advantage to the cell for loss, retention or transfer of organellar genes to the nuclear genome uncovers a novel process that has helped shape the content and evolution of eukaryotic genomes.

## Materials and Methods

### Data sources

The *Arabidopsis thaliana* genome sequence and corresponding set of representative gene models were downloaded from Phytozome V13 (Goodstein, et al. 2012). The human genome sequence and gene models from assembly version GRCh38.p13 (GCA_000001405.28), the *Bartonella henselae* genome sequence and gene models from assembly version ASM4670v1, the *Microcystis aeruginosa* NIES-843 genome sequence and gene models from assembly version ASM1062v1 were each downloaded from Ensembl (Yates, et al. 2020). The *Saccharomyces cerevisiae* sequence and gene models from assembly version R64-2-1_20150113 were downloaded from the *Saccharomyces* Genome Database (Cherry, et al. 2012). Protein abundance data for all species were obtained from PAXdb v4.1 (Wang, et al. 2015).

### Constants used to evaluate the per cell ATP costs of genes and chromosomes

The ATP biosynthesis cost of nucleotides and amino acids was obtained from (Chen, et al. 2016) and (Lynch and Marinov 2015) and are provided in Supplemental Table S3. *The Homo sapiens* mitochondrial genome copy number of 5000 was obtained from (Cole 2016). The *Saccharomyces cerevisiae* mitochondrial genome copy number of 200 was obtained from (Miyakawa 2017). The *Arabidopsis thaliana* chloroplast genome copy number of 1500 was obtained from (Zoschke, et al. 2007) and the *Arabidopsis thaliana* mitochondrial genome copy number of 100 was obtained from (Cole 2016).

For genes in nuclear chromosomes the cost of DNA was calculated to include the cost of nucleosomes with one histone octamer comprising two copies each of the histone proteins H2A, H2B, H3, and H4 every 180bp (147bp for the two turns of DNA around the histone octamer and 33bp for the spacer) (Lynch and Marinov 2015). For organellar chromosomes there are no histones/nucleosomes and thus the biosynthetic cost of genes in organellar chromosomes was calculated as cost of the DNA divided by the number of genes on the chromosome (Supplemental Table S4). Although there are no histone protein equivalents in that organellar genomes, it should be noted that there are some nuclear encoded proteins that are known to bind mitochondrial or chloroplast DNA. The costs associated with these proteins have not been included here as their function in packaging DNA is unknown and their density within the organellar genome is also unknown and it is thus difficult to estimate their required abundance. However, inclusion of the production and import costs of these proteins would further increase the cost of encoding a gene in the organellar genome and would accentuate the differences shown in this study.

The average gene length used for the simulation study in Figure 2 was obtained by computing the average gene length across the two bacterial genomes used in this study, *Bartonella henselae* ASM4670v1 and Microcystis *aeruginosa* NIES-843.

### Calculating protein import costs

Although the molecular mechanisms of mitochondrial and chloroplast protein import differ (Soll and Schleiff 2004; Jarvis 2008; Wiedemann and Pfanner 2017) they share many commonalities including the requirement for energy in the form of nucleoside triphosphate hydrolysis (Schatz and Dobberstein 1996). The energetic cost of mitochondrial or chloroplast protein import is difficult to measure directly, and accordingly estimates vary over two orders of magnitude from ∼0.05 ATP per amino acid to 5 ATP per amino acid (Mokranjac and Neupert 2008; Shi and Theg 2013; Backes and Herrmann 2017). Thus, for the purposes of this study the full range of estimates was considered in all simulations when evaluating the import cost of organellar targeted proteins encoded by nuclear genes.

The cost of the biosynthesis of the protein import machinery (i.e. the TOC/TIC or TOM/TIM complexes, Supplemental Table S5) was also included in the per protein import costs calculated in this study. For *Arabidopsis thaliana,* if the total ATP biosynthesis cost of all TOC/TIC complex proteins in the cell (i.e. the full biosynthesis cost of all the amino acids of all the proteins at their measured abundance in the cell) is distributed equally among all of the proteins that are imported into the chloroplast then it would add an additional 0.2 ATP per residue imported (Supplemental Table S6). Similarly, if the total ATP biosynthesis cost of all TOM/TIM proteins in the cell in *Homo sapiens*, *Saccharomyces cerevisiae* and *Arabidopsis thaliana* is distributed equally among all of the proteins that are imported into the mitochondrion in those species then it would add an additional 0.2 ATP, 0.7 ATP, and 0.2 ATP per residue imported, respectively (Supplemental Table S6). In all cases the proteins that were predicted to be imported into the organelle were identified using TargetP-2.0 (Almagro Armenteros, et al. 2019) and protein abundance was calculated using measured protein abundance estimates for each species obtained from PAXdb 4.0 (Wang, et al. 2015), assuming a total cell protein content of 1×10^9^ proteins for a human cell, 1×10^7^ proteins for a yeast cell and 2.5 × 10^10^ proteins for an *Arabidopsis thaliana* cell. As we modelled ATP import costs from 0.05 ATP to 50 ATP per-residue the cost of the import machinery was considered to be included within the bounds considered in this analysis.

### Evaluating the proportion of the total proteome invested in organelles

To provide estimates of the fraction of cellular protein resources invested in organellar proteomes the complete predicted proteomes and corresponding protein abundances were quantified. Organellar targeting was predicted using TargetP-2.0 (Almagro Armenteros, et al. 2019) and protein abundance estimates obtained from PAXdb 4.0 (Wang, et al. 2015). The proportion of cellular resources are provided in Supplemental Table S1 and were used to provide the indicative regions or parameter space occupied by metazoa, yeast and plants shown on Figure 2B and C. Specifically, ∼5% of total cellular protein is contained within mitochondria in *H. sapiens*, *S. cerevisiae* and *A. thaliana* and ∼50% of total cellular protein is contained within chloroplasts in *A. thaliana*.

### Calculating the free energy of endosymbiotic gene transfer

The free energy of endosymbiotic gene transfer (*ΔE_EGT_*) is here defined as the difference in energy cost to the cell to encode a given gene in the organellar genome and the cost to encode the same gene in the nuclear genome and import the requisite amount of gene product into to the organelle. *ΔE_EGT_* is evaluated as the difference in ATP biosynthesis cost required to encode a gene (*ΔD*) in the endosymbiont genome (*D_end_*) and the nuclear genome (*D_nuc_*) minus the difference in ATP biosynthesis cost required to produce the protein (*ΔP*) in the organelle (*P_end_*) vs in the cytosol (*P_cyt_*) and ATP cost to import the protein into the organelle (*P_import_*). Such that

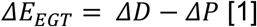

Where

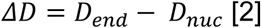

And

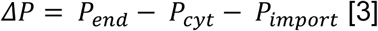

Thus, *ΔE_EGT_* can be positive or negative depending on the cost associated with each parameter. The energetic cost of producing a protein in the endosymbiont and in the cytosol are assumed to be equal and thus

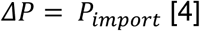

It should be noted here that although the *P_end_* and *P_cyt_* are assumed to be equal for the majority of calculations an analysis was conducted wherein an inefficient protein import system was assumed such that 50% of protein failed to be imported and thus must be turned over (Supplemental Figure S2). Even under these conditions it is still energetically favourable to encode organellar genes in the nuclear genome for realistic estimates of cell sizes and investment in organelles.

*P_import_* is evaluated as the product of the product of the length of the amino acid sequence (*L_prot_*), the ATP cost of importing a single residue from the contiguous polypeptide chain of that protein (*C_import_*), the number of copies of that protein contained within the cell that must be imported (*N_p_*) such that

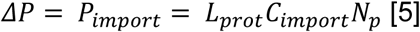

Measured estimates of *C_import_* range from ∼0.05 ATP per amino acid to 5 ATP per amino acid (Mokranjac and Neupert 2008; Shi and Theg 2013; Backes and Herrmann 2017). For the purposes of this study we used these measured ranges and also modelled a *C_import_* up to 10 times higher than any measured estimate i.e. from 0.05 ATP to 50 ATP.

Both *D_end_* and *D_nuc_* are evaluated as the product of the ATP biosynthesis cost of the double stranded DNA (*A_DNA_*) that comprises the gene under consideration and the copy number (*C*) of the genome in the cell such that

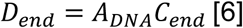

And

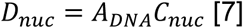

Such that

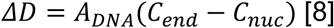

Where *C_end_* and *C_nuc_* are the per-cell copy number of the endosymbiont and nuclear genomes respectively and the ATP biosynthesis cost for the complete biosynthesis of an A:T base pair and a G:C base pair are 40.55 ATP and 40.14 ATP respectively (Chen, et al. 2016). Thus

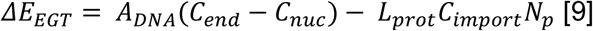

Where positive values of *ΔE_EGT_* correspond to genes for which it is more energetically favourable to be encoded in the nuclear genome, and negative values correspond to genes for which it is more energetically favourable to be encoded in the endosymbiont genome.

### Simulating endosymbiotic gene transfer of mitochondrial and chloroplast genes

The complete genomes with measured protein abundances for an alphaproteobacterium (*Bartonella henselae*) and a cyanobacterium (*Microcystis aeruginosa*) were selected to sever as models for an ancestral mitochondrion and cyanobacterium, respectively. To account for uncertainty in the size and complexity of the ancestral pre-mitochondrial and pre-chloroplast host cells, a range of potential ancestral cells was considered to be engulfed by a range of different host cells with protein contents representative of the diversity of extant eukaryotic cells (Milo 2013). Specifically, the size of the host cell ranged from a small unicellular yeast-like cell (10^7^ proteins), to a medium sized unicellular algal-like cell (10^8^ proteins) to a typical metazoan/plant cell (10^9^ proteins). Each of these host cell types was then considered to allocate a realistic range of total cellular protein to mitochondria/chloroplasts typical of eukaryotic cells (i.e. ∼2% for yeast (Uchida, et al. 2011), ∼20% for metazoan cells (David 1977) and ∼50% of the non-vacuolar volume of plant cells (Winter, et al. 1994)). It is not important whether the organellar fraction of the cell is composed of a single large organelle or multiple smaller organelles as all costs, abundances, and copy numbers are evaluated at a per-cell level. For each simulated cell, *ΔE_EGT_* was evaluated for each gene in the endosymbiont genome using real protein abundance data (Wang, et al. 2015) for a realistic range of endosymbiont genome copy numbers using equation 9. In all cases the host cell was assumed to be diploid. The simulations were repeated for three different per-residue protein import costs (0.05 ATP, 2 ATP, and 5 ATP per residue respectively). The number of genes where *ΔE_EGT_* was positive was recorded as these genes comprise the cohort that are energetically favourable to be encoded in the nuclear genome. All calculated values for *ΔE_EGT_* for both the model organisms are provided in Supplemental Table S2.

### Estimating the strength of selection acting on endosymbiotic gene transfer

To model the proportion of energy that would be saved by an individual endosymbiotic gene transfer event a number of assumptions were made. It was assumed that the ancestral host cell had a cell size that is within the range of extant eukaryotes (i.e. between 1 × 10^7^ proteins per cell and 1 × 10^9^ proteins per cell). It was assumed that the endosymbiont occupied a fraction of the total cell proteome that is within the range exhibited by most eukaryotes today (2% to 50% of total cellular protein is located within the endosymbiont under consideration). It was assumed that endosymbiont genome copy number ranged between 1 copy per cell (as it most likely started out with a single copy) and 10,000 copies per cell.

We assumed an ancestral host cell with a 24-hour doubling time such that all genomes and proteins are produced in the required abundance every 24-hour period. As previously defined (Lynch and Marinov 2015), the energy required for cell growth was modelled as

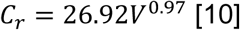

In addition, all cells, irrespective of whether they are bacterial or eukaryotic, consume ATP (*C_m_*) in proportion to their cell volume (*V*) at approximately the rate of

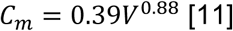

where *C_m_* is in units of 10^9^ molecules of ATP cell^-1^ hour^-1^, and V is in units of µm^3^ (Lynch and Marinov 2015). Thus, the total energy (*E_R_*) needed to replicate a cell was considered to be

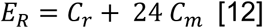

The proportional energetic advantage or disadvantage (*E_A/D_*) to the host cell from the endosymbiotic gene transfer of a given gene is evaluated as the free energy of endosymbiotic gene transfer divided by the total amount of energy consumed by the cell during its 24-hour life cycle.

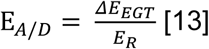

Given that *E_A/D_* describes the proportional energetic advantage or disadvantage a cell has from a given endosymbiotic gene transfer event *E_A/D_* can be used directly as selection coefficient (s) to evaluate the strength of selection acting on the endosymbiotic gene transfer of a given gene. Such that

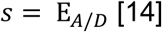

As *ΔE_EGT_* can be positive or negative as described above, s is therefore also positive or negative depending on endosymbiont genome copy number, endosymbiont fraction, host cell protein content, the abundance of the protein that must be imported and the ATP cost of protein import. When s is less than zero the absolute value of *s* is taken to be the selection coefficient for retention of a gene in the endosymbiont genome (*S_R_*), when *s* is greater than 0 the value of *s* is taken to be the selection coefficient for endosymbiotic gene transfer to the nucleus (*S_EGT_*). All calculated values for *s* for both the model alphaproteobacterium (*Bartonella henselae*) and cyanobacterium (*Microcystis aeruginosa*) are provided in Supplemental Table S1.

It should be noted here that variation in the doubling time will have a direct effect on the estimate of the selection coefficients. Decreasing cell doubling time 10-fold increases the values of *s* by a factor of 10 and *vice versa*, such that cells with shorter doubling times would experience stronger selection on the free energy of endosymbiotic gene transfer and cells with longer doubling times would experience weaker selection.

Similarly, the effect of protein turnover was not included as estimates for protein turn over were not available for each protein considered in these analyses. However, the effect of protein turnover is to increase the total amount of protein that must be produced within the life cycle of the cell. Thus, for the purposes of this analysis can be considered equivalent to increasing the cellular investment in organelles.

### Estimating time to fixation

Fixation times for endosymbiotic gene transfer events for a range of observed selection coefficients from 1 × 10^-5^ to 1 × 10^-2^ were estimated using a Wright–Fisher model with selection and drift (Fisher 1930; Wright 1931) implemented in a simple evolutionary dynamics simulation (Niklaus and Kelly 2018). The effective population size for these simulations was set as 1 × 10^7^, as is representative of unicellular eukaryotes (Lynch and Conery 2003) and multicellularity in eukaryotes is not thought to have evolved until after the endosymbiosis of either the mitochondrion or the chloroplast.

## Acknowledgements and funding sources

This work was funded by the Royal Society and the European Union’s Horizon 2020 research and innovation program under grant agreement number 637765. SK would like to thank Thomas A. Richards, David M. Emms, and John M. Archibald for their comments on the manuscript.

## Data availability

All of the data required to conduct this analysis was obtained from public repositories as outlined in the materials and methods.

## References

Ahmadinejad N, Dagan T, Gruenheit N, Martin W, Gabaldón T. 2010. Evolution of spliceosomal introns following endosymbiotic gene transfer. BMC Evol Biol 10:57.

Allen JF, Raven JA. 1996. Free-radical-induced mutation vs redox regulation: costs and benefits of genes in organelles. J Mol Evol 42:482–492.

Almagro Armenteros JJ, Salvatore M, Emanuelsson O, Winther O, von Heijne G, Elofsson A, Nielsen H. 2019. Detecting sequence signals in targeting peptides using deep learning. Life Sci Alliance 2.

Archibald John M. 2015a. Endosymbiosis and Eukaryotic Cell Evolution. Current Biology 25:R911–R921.

Archibald JM. 2015b. Genomic perspectives on the birth and spread of plastids. Proceedings of the National Academy of Sciences 112:10147–10153.

Backes S, Herrmann JM. 2017. Protein Translocation into the Intermembrane Space and Matrix of Mitochondria: Mechanisms and Driving Forces. Frontiers in Molecular Biosciences 4.

Bar-On YM, Phillips R, Milo R. 2018. The biomass distribution on Earth. Proceedings of the National Academy of Sciences 115:6506–6511.

Bendich AJ. 1987. Why do chloroplasts and mitochondria contain so many copies of their genome? Bioessays 6:279–282.

Bersaglieri T, Sabeti PC, Patterson N, Vanderploeg T, Schaffner SF, Drake JA, Rhodes M, Reich DE, Hirschhorn JN. 2004. Genetic signatures of strong recent positive selection at the lactase gene. Am J Hum Genet 74:1111–1120.

Boisvert FM, Ahmad Y, Gierliński M, Charrière F, Lamont D, Scott M, Barton G, Lamond AI. 2012. A quantitative spatial proteomics analysis of proteome turnover in human cells. Mol Cell Proteomics 11:M111.011429.

Booth A, Doolittle WF. 2015a. Eukaryogenesis, how special really? Proceedings of the National Academy of Sciences 112:10278–10285.

Booth A, Doolittle WF. 2015b. Reply to Lane and Martin: Being and becoming eukaryotes. Proceedings of the National Academy of Sciences 112:E4824–E4824.

Brown JR. 2003. Ancient horizontal gene transfer. Nature Reviews Genetics 4:121–132.

Calvo SE, Mootha VK. 2010. The mitochondrial proteome and human disease. Annu Rev Genomics Hum Genet 11:25–44.

Chen W-H, Lu G, Bork P, Hu S, Lercher MJ. 2016. Energy efficiency trade-offs drive nucleotide usage in transcribed regions. Nature Communications 7:11334.

Cherry JM, Hong EL, Amundsen C, Balakrishnan R, Binkley G, Chan ET, Christie KR, Costanzo MC, Dwight SS, Engel SR, et al. 2012. Saccharomyces Genome Database: the genomics resource of budding yeast. Nucleic Acids Res 40:D700–705.

Cole LW. 2016. The Evolution of Per-cell Organelle Number. Front Cell Dev Biol 4:85.

Dagan T, Roettger M, Stucken K, Landan G, Koch R, Major P, Gould SB, Goremykin VV, Rippka R, Tandeau de Marsac N, et al. 2013. Genomes of Stigonematalean cyanobacteria (subsection V) and the evolution of oxygenic photosynthesis from prokaryotes to plastids. Genome Biol Evol 5:31–44.

Daley DO, Whelan J. 2005. Why genes persist in organelle genomes. Genome Biology 6:110.

David H. 1977. Quantitative Ultrastructural Data of Animal and Human Cells: Gustav Fischer.

Deusch O, Landan G, Roettger M, Gruenheit N, Kowallik KV, Allen JF, Martin W, Dagan T. 2008. Genes of Cyanobacterial Origin in Plant Nuclear Genomes Point to a Heterocyst-Forming Plastid Ancestor. Molecular Biology and Evolution 25:748–761.

Ding WX, Yin XM. 2012. Mitophagy: mechanisms, pathophysiological roles, and analysis. Biol Chem 393:547–564.

Doolittle WF. 1998. You are what you eat: a gene transfer ratchet could account for bacterial genes in eukaryotic nuclear genomes. Trends Genet 14:307–311.

Drouin G, Daoud H, Xia J. 2008. Relative rates of synonymous substitutions in the mitochondrial, chloroplast and nuclear genomes of seed plants. Mol Phylogenet Evol 49:827–831.

Ferro M, Brugière S, Salvi D, Seigneurin-Berny D, Court M, Moyet L, Ramus C, Miras S, Mellal M, Le Gall S, et al. 2010. AT_CHLORO, a comprehensive chloroplast proteome database with subplastidial localization and curated information on envelope proteins. Mol Cell Proteomics 9:1063–1084.

Fisher RAS. 1930. The genetical theory of natural selection. Oxford: Clarendon Press.

Gallaher SD, Craig RJ, Ganesan I, Purvine SO, McCorkle SR, Grimwood J, Strenkert D, Davidi L, Roth MS, Jeffers TL, et al. 2021. Widespread polycistronic gene expression in green algae. Proceedings of the National Academy of Sciences 118:e2017714118.

Gawron D, Ndah E, Gevaert K, Van Damme P. 2016. Positional proteomics reveals differences in N-terminal proteoform stability. Mol Syst Biol 12:858.

Goodstein DM, Shu S, Howson R, Neupane R, Hayes RD, Fazo J, Mitros T, Dirks W, Hellsten U, Putnam N, et al. 2012. Phytozome: a comparative platform for green plant genomics. Nucleic Acids Res 40:D1178–1186.

Gordon SP, Tseng E, Salamov A, Zhang J, Meng X, Zhao Z, Kang D, Underwood J, Grigoriev IV, Figueroa M, et al. 2015. Widespread Polycistronic Transcripts in Fungi Revealed by Single-Molecule mRNA Sequencing. PLoS One 10:e0132628.

Gray MW, Burger G, Lang BF. 1999. Mitochondrial evolution. Science 283:1476–1481.

Green BR. 2011. Chloroplast genomes of photosynthetic eukaryotes. Plant J 66:34–44.

Grisdale CJ, Smith DR, Archibald JM. 2019. Relative Mutation Rates in Nucleomorph-Bearing Algae. Genome Biology and Evolution 11:1045–1053.

Guiliano DB, Blaxter ML. 2006. Operon conservation and the evolution of trans-splicing in the phylum Nematoda. PLoS Genet 2:e198.

Hartl DL, Moriyama EN, Sawyer SA. 1994. Selection intensity for codon bias. Genetics 138:227–234.

Herrmann R. 1997. Eukaryotism, towards a new interpretation. In. Eukaryotism and symbiosis: Springer. p. 73–118.

Husnik F, Nikoh N, Koga R, Ross L, Duncan RP, Fujie M, Tanaka M, Satoh N, Bachtrog D, Wilson AC, et al. 2013. Horizontal gene transfer from diverse bacteria to an insect genome enables a tripartite nested mealybug symbiosis. Cell 153:1567–1578.

Jarvis P. 2008. Targeting of nucleus-encoded proteins to chloroplasts in plants. New Phytologist 179:257–285.

Khakhlova O, Bock R. 2006. Elimination of deleterious mutations in plastid genomes by gene conversion. The Plant Journal 46:85–94.

Lane N. 2014. Bioenergetic constraints on the evolution of complex life. Cold Spring Harb Perspect Biol 6:a015982.

Lane N, Martin W. 2010. The energetics of genome complexity. Nature 467:929–934.

Lane N, Martin WF. 2015. Eukaryotes really are special, and mitochondria are why. Proceedings of the National Academy of Sciences 112:E4823–E4823.

Lynch M. 1997. Mutation accumulation in nuclear, organelle, and prokaryotic transfer RNA genes. Mol Biol Evol 14:914–925.

Lynch M. 1996. Mutation accumulation in transfer RNAs: molecular evidence for Muller’s ratchet in mitochondrial genomes. Mol Biol Evol 13:209–220.

Lynch M, Conery JS. 2003. The origins of genome complexity. Science 302:1401–1404.

Lynch M, Koskella B, Schaack S. 2006. Mutation Pressure and the Evolution of Organelle Genomic Architecture. Science 311:1727–1730.

Lynch M, Lynch PSTSM, Walsh B. 2007. The Origins of Genome Architecture: Oxford University Press, Incorporated.

Lynch M, Marinov GK. 2015. The bioenergetic costs of a gene. Proceedings of the National Academy of Sciences 112:15690–15695.

Lynch M, Marinov GK. 2017. Membranes, energetics, and evolution across the prokaryote-eukaryote divide. eLife 6:e20437.

Lynch M, Marinov GK. 2018. Response to Martin and colleagues: mitochondria do not boost the bioenergetic capacity of eukaryotic cells. Biology Direct 13:26.

Martin-Perez M, Villén J. 2017. Determinants and Regulation of Protein Turnover in Yeast. Cell Syst 5:283–294.e285.

Martin W, Herrmann RG. 1998. Gene Transfer from Organelles to the Nucleus: How Much, What Happens, and Why? Plant Physiology 118:9–17.

Martin W, Kowallik K. 1999. Annotated English translation of Mereschkowsky’s 1905 paper ‘Über Natur und Ursprung der Chromatophoren imPflanzenreiche’. European Journal of Phycology 34:287–295.

Martin W, Müller M. 1998. The hydrogen hypothesis for the first eukaryote. Nature 392:37–41.

Martin W, Rujan T, Richly E, Hansen A, Cornelsen S, Lins T, Leister D, Stoebe B, Hasegawa M, Penny D. 2002. Evolutionary analysis of *Arabidopsis*, cyanobacterial, and chloroplast genomes reveals plastid phylogeny and thousands of cyanobacterial genes in the nucleus. Proceedings of the National Academy of Sciences 99:12246–12251.

Martin WF, Garg S, Zimorski V. 2015. Endosymbiotic theories for eukaryote origin. Philos Trans R Soc Lond B Biol Sci 370:20140330.

McCutcheon JP, Moran NA. 2012. Extreme genome reduction in symbiotic bacteria. Nature Reviews Microbiology 10:13–26.

Michaeli S. 2011. Trans-splicing in trypanosomes: machinery and its impact on the parasite transcriptome. Future Microbiol 6:459–474.

Milo R. 2013. What is the total number of protein molecules per cell volume? A call to rethink some published values. Bioessays 35:1050–1055.

Miyakawa I. 2017. Organization and dynamics of yeast mitochondrial nucleoids. Proc Jpn Acad Ser B Phys Biol Sci 93:339–359.

Mokranjac D, Neupert W. 2008. Energetics of protein translocation into mitochondria. Biochimica et Biophysica Acta (BBA) - Bioenergetics 1777:758–762.

Muller HJ. 1964. The relation of recombination to mutational advance. Mutation Research/Fundamental and Molecular Mechanisms of Mutagenesis 1:2–9.

Nakayama T, Ishida K. 2009. Another acquisition of a primary photosynthetic organelle is underway in Paulinella chromatophora. Curr Biol 19:R284–285.

Neiman M, Taylor DR. 2009. The causes of mutation accumulation in mitochondrial genomes. Proceedings of the Royal Society B: Biological Sciences 276:1201–1209.

Niklaus M, Kelly S. 2018. The molecular evolution of C4 photosynthesis: opportunities for understanding and improving the world’s most productive plants. Journal of Experimental Botany 70:795–804.

Nowack EC, Price DC, Bhattacharya D, Singer A, Melkonian M, Grossman AR. 2016. Gene transfers from diverse bacteria compensate for reductive genome evolution in the chromatophore of Paulinella chromatophora. Proc Natl Acad Sci U S A 113:12214–12219.

Nowack ECM, Vogel H, Groth M, Grossman AR, Melkonian M, Glöckner G. 2010. Endosymbiotic Gene Transfer and Transcriptional Regulation of Transferred Genes in Paulinella chromatophora. Molecular Biology and Evolution 28:407–422.

Nowack ECM, Weber APM. 2018. Genomics-Informed Insights into Endosymbiotic Organelle Evolution in Photosynthetic Eukaryotes. Annual Review of Plant Biology 69:51–84.

Pittis AA, Gabaldon T. 2016. Late acquisition of mitochondria by a host with chimaeric prokaryotic ancestry. Nature 531:101–104.

Reyes-Prieto A. 2015. The basic genetic toolkit to move in with your photosynthetic partner. Frontiers in Ecology and Evolution 3.

Reyes-Prieto A, Hackett JD, Soares MB, Bonaldo MF, Bhattacharya D. 2006. Cyanobacterial Contribution to Algal Nuclear Genomes Is Primarily Limited to Plastid Functions. Current Biology 16:2320–2325.

Reyes-Prieto A, Yoon HS, Moustafa A, Yang EC, Andersen RA, Boo SM, Nakayama T, Ishida K, Bhattacharya D. 2010. Differential gene retention in plastids of common recent origin. Mol Biol Evol 27:1530–1537.

Roger AJ, Muñoz-Gómez SA, Kamikawa R. 2017. The Origin and Diversification of Mitochondria. Curr Biol 27:R1177–r1192.

Rogozin IB, Carmel L, Csuros M, Koonin EV. 2012. Origin and evolution of spliceosomal introns. Biol Direct 7:11.

Schatz G, Dobberstein B. 1996. Common Principles of Protein Translocation Across Membranes. Science 271:1519–1526.

Shi L-X, Theg SM. 2013. Energetic cost of protein import across the envelope membranes of chloroplasts. Proceedings of the National Academy of Sciences 110:930–935.

Shokolenko I, Venediktova N, Bochkareva A, Wilson GL, Alexeyev MF. 2009. Oxidative stress induces degradation of mitochondrial DNA. Nucleic Acids Res 37:2539–2548.

Singer A, Poschmann G, Mühlich C, Valadez-Cano C, Hänsch S, Hüren V, Rensing SA, Stühler K, Nowack ECM. 2017. Massive Protein Import into the Early-Evolutionary-Stage Photosynthetic Organelle of the Amoeba Paulinella chromatophora. Curr Biol 27:2763–2773.e2765.

Smith DR. 2015. Mutation rates in plastid genomes: they are lower than you might think. Genome Biol Evol 7:1227–1234.

Smith DR. 2016. The mutational hazard hypothesis of organelle genome evolution: 10 years on. Molecular Ecology 25:3769–3775.

Smith DR, Keeling PJ. 2015. Mitochondrial and plastid genome architecture: Reoccurring themes, but significant differences at the extremes. Proceedings of the National Academy of Sciences 112:10177–10184.

Soll J, Schleiff E. 2004. Protein import into chloroplasts. Nat Rev Mol Cell Biol 5:198–208.

Speijer D, Hammond M, Lukeš J. 2020. Comparing Early Eukaryotic Integration of Mitochondria and Chloroplasts in the Light of Internal ROS Challenges: Timing is of the Essence. mBio 11:e00955–00920.

Thiergart T, Landan G, Schenk M, Dagan T, Martin WF. 2012. An evolutionary network of genes present in the eukaryote common ancestor polls genomes on eukaryotic and mitochondrial origin. Genome Biol Evol 4:466–485.

Timmis JN, Ayliffe MA, Huang CY, Martin W. 2004. Endosymbiotic gene transfer: organelle genomes forge eukaryotic chromosomes. Nat Rev Genet 5:123–135.

Uchida M, Sun Y, McDermott G, Knoechel C, Le Gros MA, Parkinson D, Drubin DG, Larabell CA. 2011. Quantitative analysis of yeast internal architecture using soft X-ray tomography. Yeast 28:227–236.

Wang M, Herrmann CJ, Simonovic M, Szklarczyk D, von Mering C. 2015. Version 4.0 of PaxDb: Protein abundance data, integrated across model organisms, tissues, and cell-lines. Proteomics 15:3163–3168.

Wiedemann N, Pfanner N. 2017. Mitochondrial Machineries for Protein Import and Assembly. Annual Review of Biochemistry 86:685–714.

Winter H, Robinson DG, Heldt HW. 1994. Subcellular volumes and metabolite concentrations in spinach leaves. Planta 193:530–535.

Wolfe KH, Li WH, Sharp PM. 1987. Rates of nucleotide substitution vary greatly among plant mitochondrial, chloroplast, and nuclear DNAs. Proceedings of the National Academy of Sciences 84:9054–9058.

Wright S. 1931. Evolution in Mendelian Populations. Genetics 16:97–159.

Yang D, Oyaizu Y, Oyaizu H, Olsen GJ, Woese CR. 1985. Mitochondrial origins. Proceedings of the National Academy of Sciences 82:4443–4447.

Yates AD, Achuthan P, Akanni W, Allen J, Allen J, Alvarez-Jarreta J, Amode MR, Armean IM, Azov AG, Bennett R, et al. 2020. Ensembl 2020. Nucleic Acids Res 48:D682–d688.

Zhuang X, Jiang L. 2019. Chloroplast Degradation: Multiple Routes Into the Vacuole. Frontiers in Plant Science 10.

Zoschke R, Liere K, Börner T. 2007. From seedling to mature plant: Arabidopsis plastidial genome copy number, RNA accumulation and transcription are differentially regulated during leaf development. The Plant Journal 50:710–722.

