## Supplemental Figures S1 - S9 for "The economics of organellar gene loss and endosymbiotic gene transfer"

### *Supplemental Figure S1*

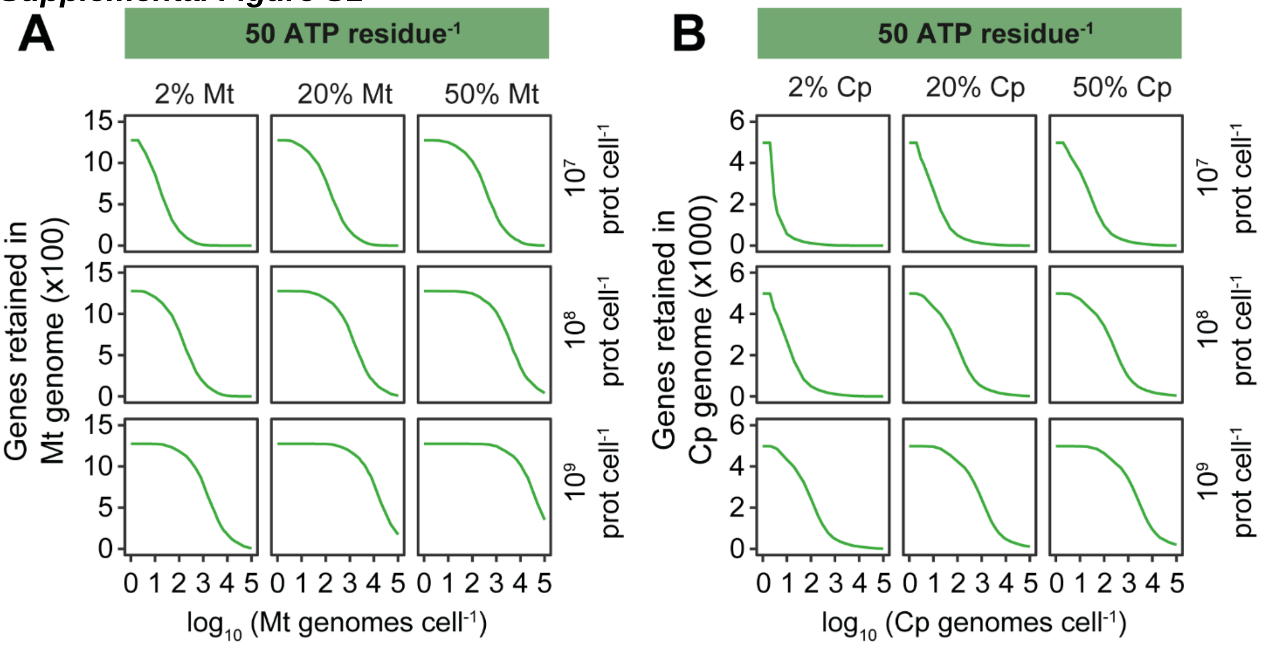


**Supplemental Figure S1.** The effect of assuming an ATP import cost 10 fold higher than the upper estimates of import cost measured in cells. **A)** The number of genes in the alphaproteobacterial (mitochondrial) genome for which it is more energetically favourable to the cell for the gene to be retained in the organellar genome. Green lines assume a per-residue protein import cost of 50 ATP per amino acid. **B**) As in A but for the cyanobacterial (chloroplast) genome.

***Supplemental Figure S2***

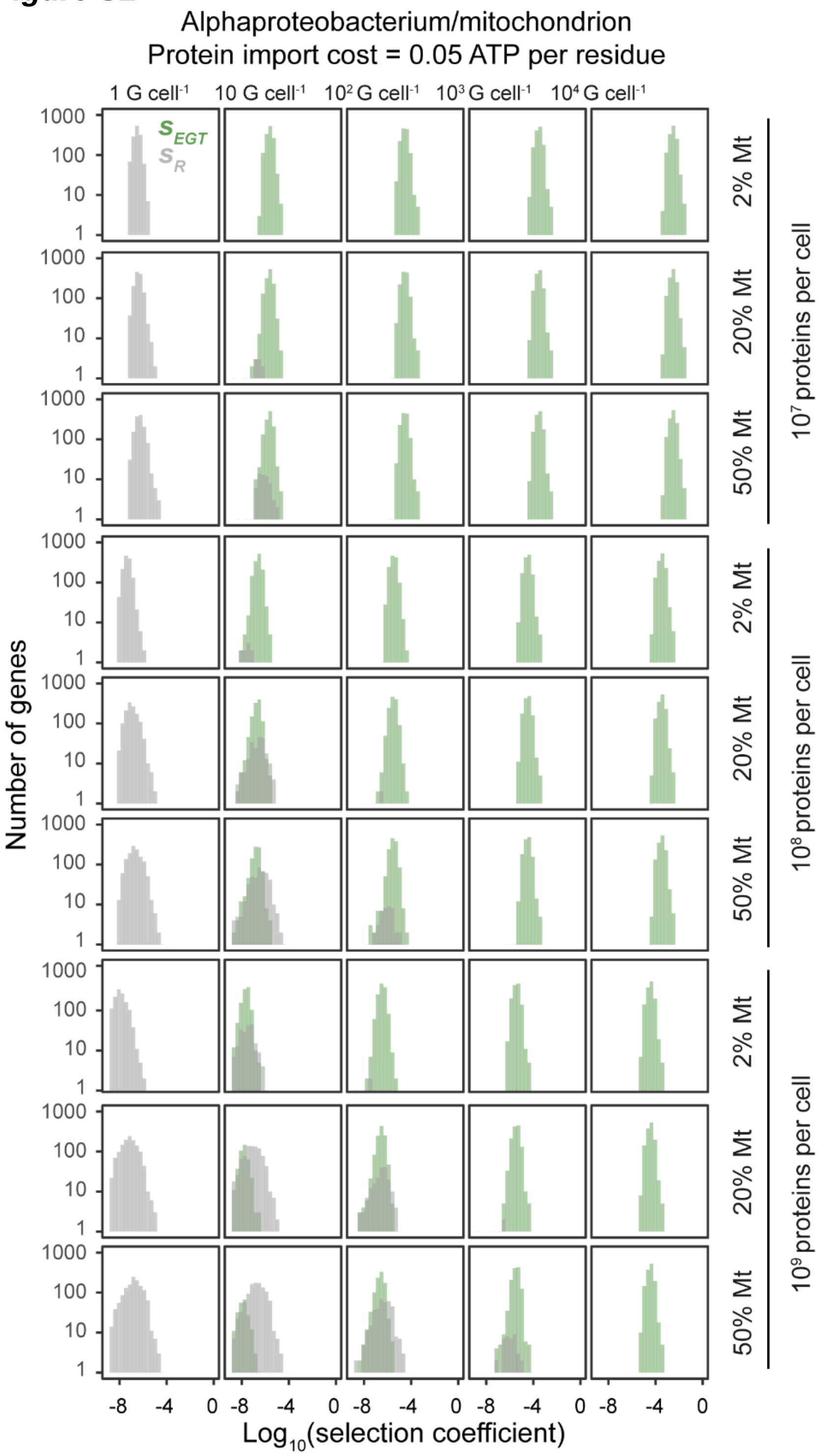


**Supplemental Figure S2** The selection coefficients for endosymbiotic gene transfer of alphaproteobacterial genes as a function of host cell size, host cell mitochondrial fraction and mitochondrial genome copy number per cell for a protein import cost of 0.05 ATP per residue. Histograms depict the selection coefficients for all genes in the endosymbiont genome. $S_R$ and $S_{EGT}$ have opposite signs (see methods), however to simplify the display and enable comparison the absolute value of the selection coefficients of each gene plotted.

*Supplemental Figure S3*

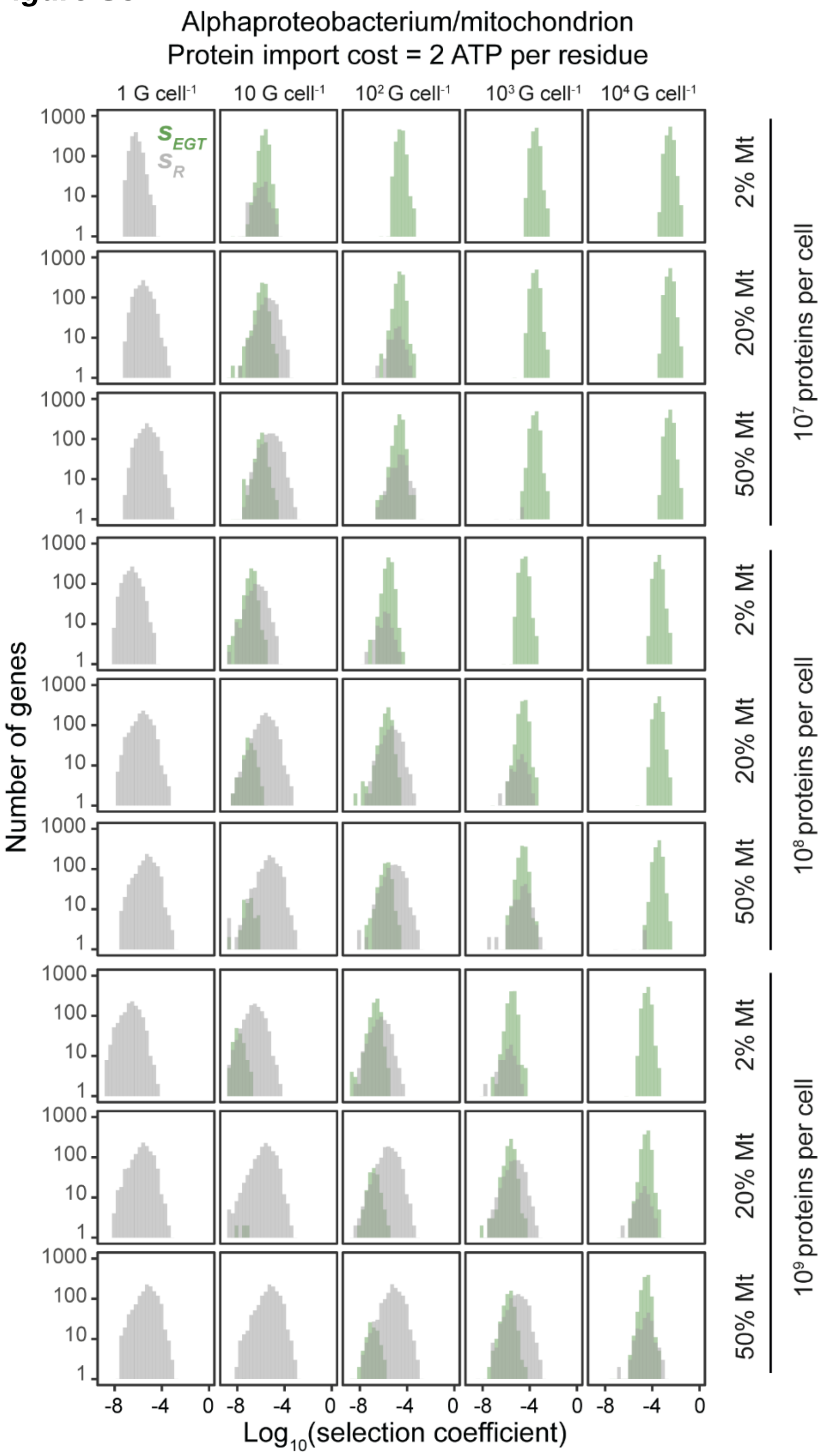


**Supplemental Figure S3.** The selection coefficients for endosymbiotic gene transfer of alphaproteobacterial genes as a function of host cell size, host cell mitochondrial fraction and mitochondrial genome copy number per cell for a protein import cost of 2 ATP per residue. Histograms depict the selection coefficients for all genes in the endosymbiont genome. $S_R$ and $S_{EGT}$ have opposite signs (see methods), however to simplify the display and enable comparison the absolute value of the selection coefficients of each gene plotted.

*Supplemental Figure S4*

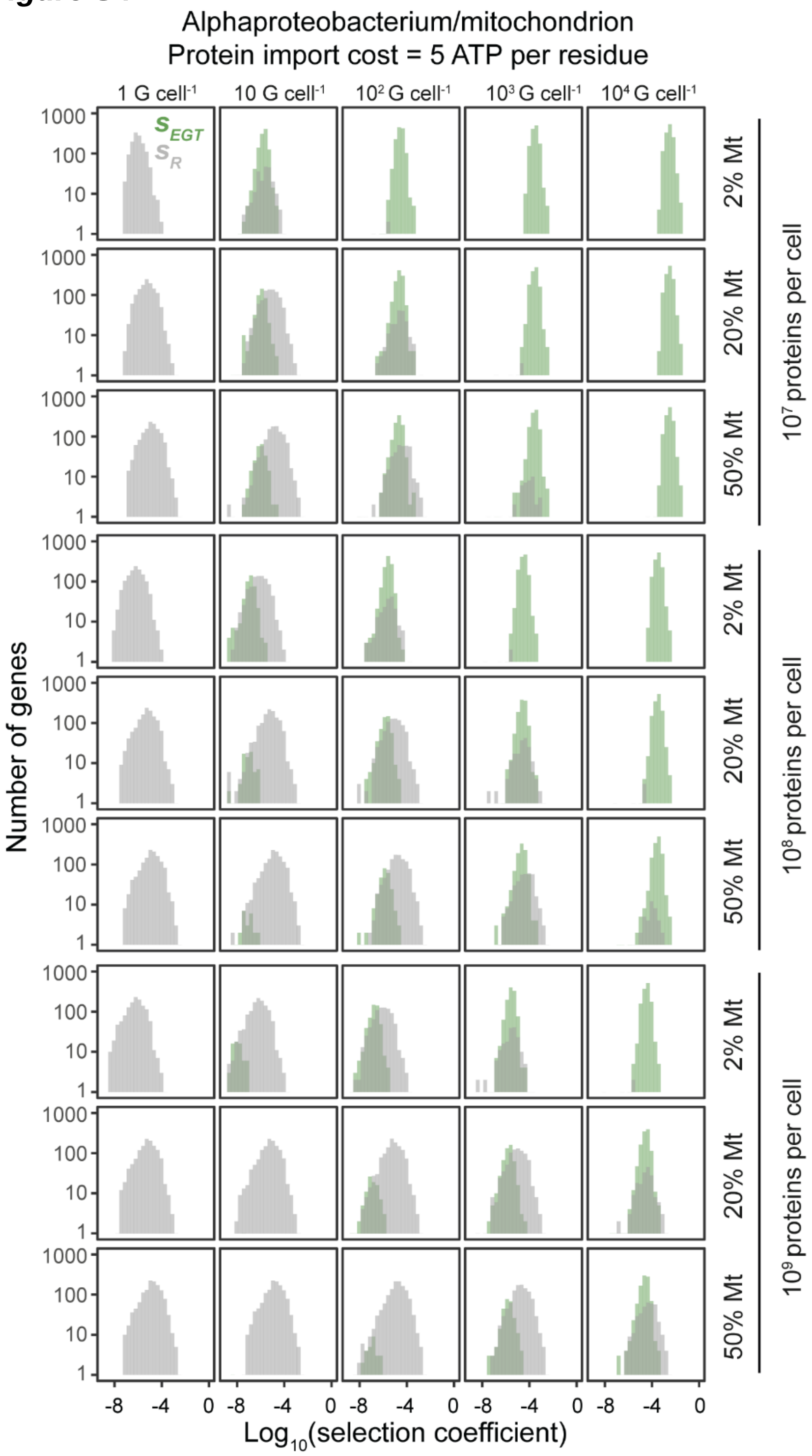


**Supplemental Figure S5.** The selection coefficients for endosymbiotic gene transfer of alphaproteobacterial genes as a function of host cell size, host cell mitochondrial fraction and mitochondrial genome copy number per cell for a protein import cost of 5 ATP per residue. Histograms depict the selection coefficients for all genes in the endosymbiont genome. $S_R$ and $S_{EGT}$ have opposite signs (see methods), however to simplify the display and enable comparison the absolute value of the selection coefficients of each gene plotted.

***Supplemental Figure S5***

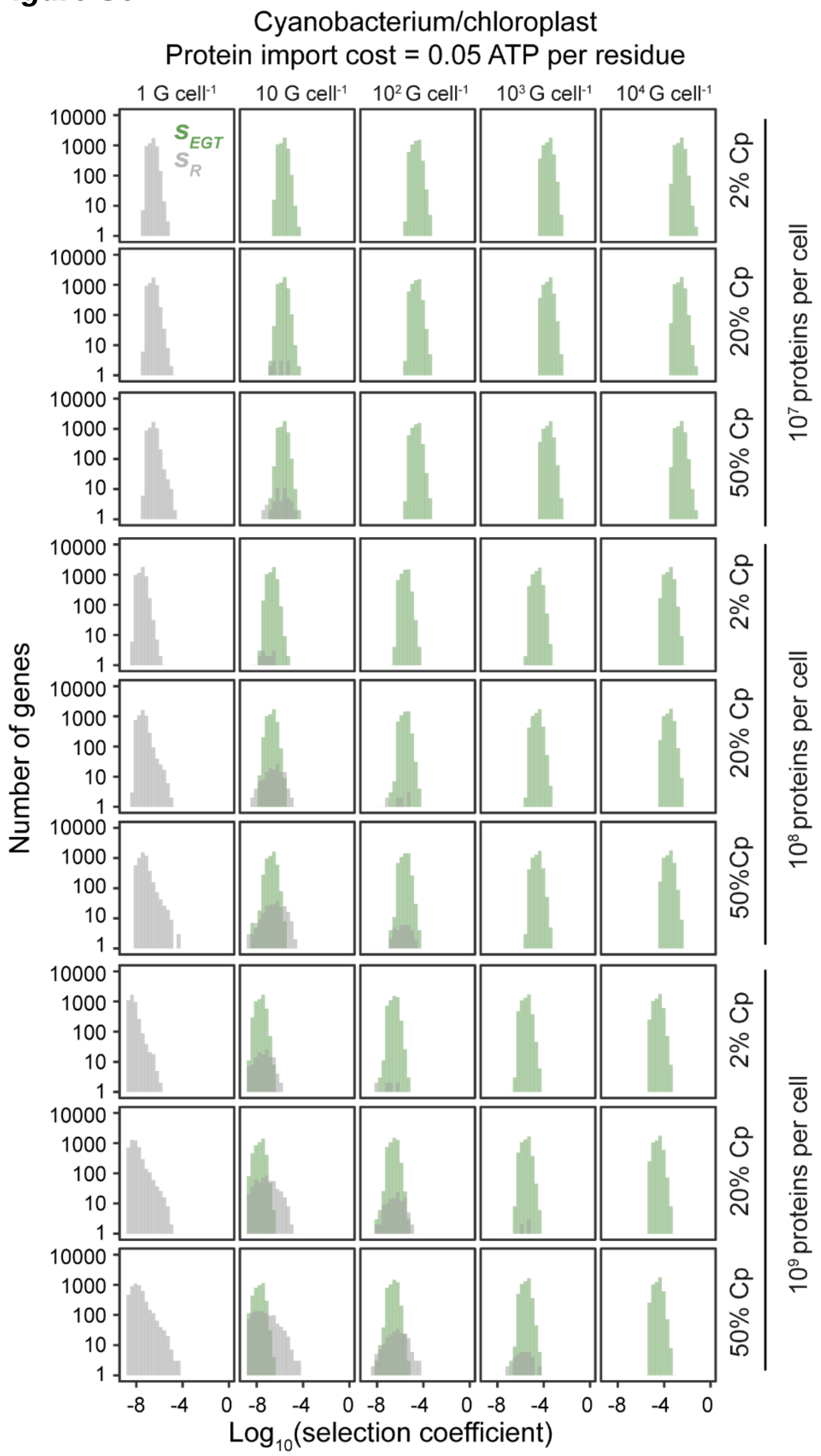


**Supplemental Figure S5.** The selection coefficients for endosymbiotic gene transfer of cyanobacterial genes as a function of host cell size, host cell chloroplast fraction and chloroplast genome copy number per cell for a protein import cost of 0.05 ATP per residue. Histograms depict the selection coefficients for all genes in the endosymbiont genome. $S_R$ and $S_{EGT}$ have opposite signs (see methods), however to simplify the display and enable comparison the absolute value of the selection coefficients of each gene plotted.

***Supplemental Figure S6***

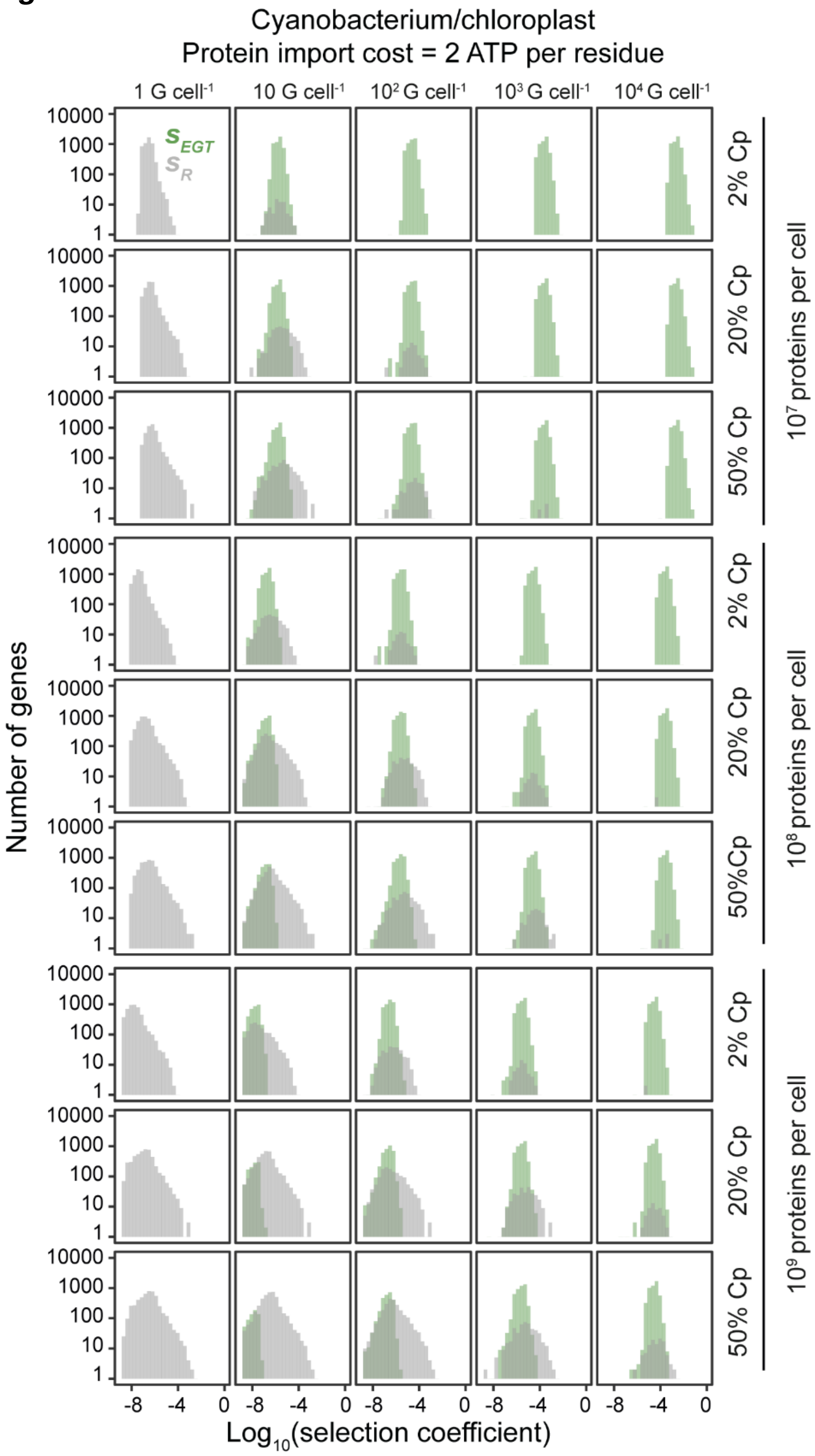


**Supplemental Figure S6.** The selection coefficients for endosymbiotic gene transfer of cyanobacterial genes as a function of host cell size, host cell chloroplast fraction and chloroplast genome copy number per cell for a protein import cost of 2 ATP per residue. Histograms depict the selection coefficients for all genes in the endosymbiont genome. $S_R$ and $S_{EGT}$ have opposite signs (see methods), however to simplify the display and enable comparison the absolute value of the selection coefficients of each gene plotted.

***Supplemental Figure S7***

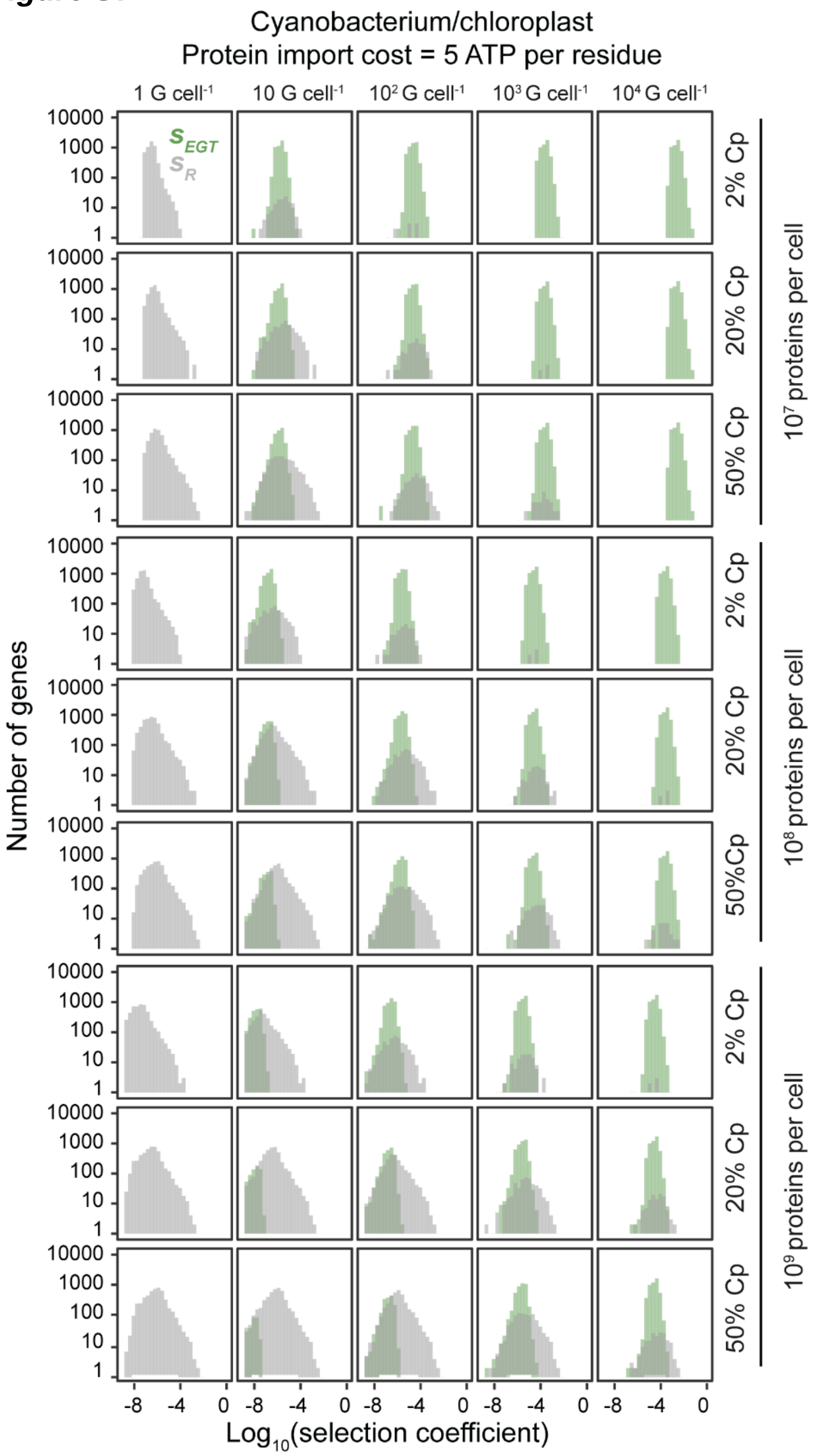


**Supplemental Figure S7.** The selection coefficients for endosymbiotic gene transfer of cyanobacterial genes as a function of host cell size, host cell chloroplast fraction and chloroplast genome copy number per cell for a protein import cost of 5 ATP per residue. Histograms depict the selection coefficients for all genes in the endosymbiont genome. $S_R$ and $S_{EGT}$ have opposite signs (see methods), however to simplify the display and enable comparison the absolute value of the selection coefficients of each gene plotted.

***Supplemental Figure S8***

| *s* | $T_{fix}$ | S.D. |
|---|---|---|
| $1 \times 10^{-1}$ | 671 | 47 |
| $1 \times 10^{-2}$ | 5830 | 576 |
| $1 \times 10^{-3}$ | 53484 | 9527 |
| $1 \times 10^{-4}$ | 479505 | 101922 |
| $1 \times 10^{-5}$ | 4263487 | 1260089 |
| $1 \times 10^{-6}$ | 38138409 | 18192217 |

**Supplemental Figure S8** Simulated fixation times ($T_{fix}$) and their standard deviations (S.D.) for a range of selection coefficients (s). Units are for $T_{fix}$ are generations.

***Supplemental Figure S9***

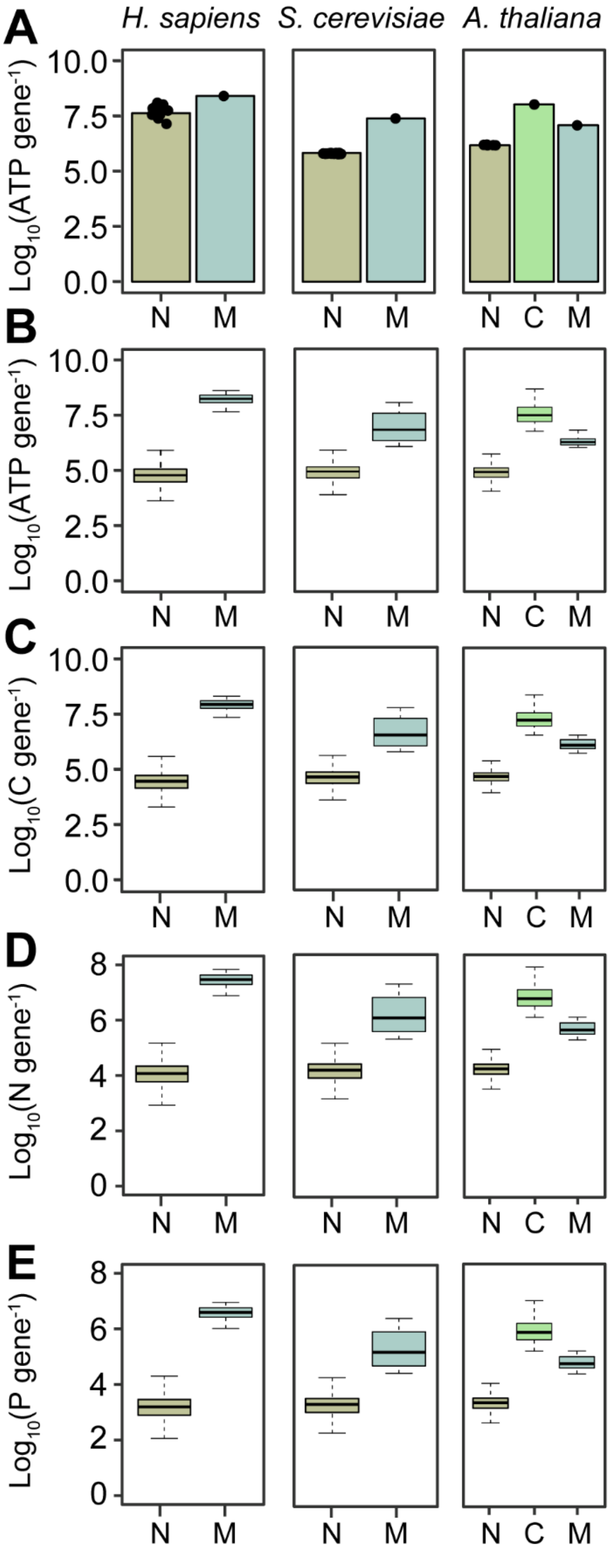


**Supplemental Figure S9.** The per-cell cost of nuclear and organellar genes in three representative eukaryotes. Part **A** and Part **B** are reproduced from the main text Figure 1 for reference. **C)** The carbon atom biosynthesis cost of nuclear (N), chloroplast (C), and mitochondrial (M) genes. **D)** The nitrogen atom biosynthesis cost of the same genes. **E)** The phosphorous atom biosynthesis cost of the same genes. Costs were computed assuming a diploid nuclear genome, a per-cell mitochondrial genome copy number of 5000, 200 and 100 for the in *H. sapiens*, *S. cerevisiae* and *A. thaliana*, respectively, and a per cell chloroplast genome copy number of 1500 in *A. thaliana*.
